# Pan-Metabolomics Repository Mapping of the Carnitine Landscape

**DOI:** 10.64898/2026.03.27.714844

**Authors:** Helena Mannochio-Russo, Patrick C. Ferreira, Kine Eide Kvitne, Abubaker Patan, Victoria Deleray, Julius Agongo, Harsha Gouda, Wilhan D. Gonçalves Nunes, Shipei Xing, Jasmine Zemlin, Martijn van Faassen, Erin R. Reilly, Imhoi Koo, Andrew D. Patterson, Shirley M. Tsunoda, Mingxun Wang, Dionicio Siegel, Lindsey A. Burnett, Pieter C. Dorrestein

## Abstract

Carnitines are a structurally diverse class of metabolites formed by conjugation of L-carnitine with fatty acids, amino acids, xenobiotics, and microbial metabolites. They play roles in transport, mitochondrial and peroxisomal metabolism, detoxification, and systemic signaling, yet their chemical diversity remains incompletely defined. We applied a pan-repository data mining strategy of LC-MS/MS data across GNPS/MassIVE, MetaboLights, and Metabolomics Workbench using MassQL diagnostic fragment ion filtering to systematically extract acylcarnitine spectra. This yielded a library of 34,222 unique MS/MS spectra representing 2,857 atomic compositions, corresponding to 3,872,050 detections. These datasets provide an MS/MS library for annotation, discovery, and contextualization of acylcarnitines, enabling identification of previously unknown carnitines, such as dihydroferulic acid conjugated carnitines and supporting future exploration of this metabolite class across host metabolism, diet, microbial activity, pharmacological exposures, and metabolic dysregulation.

## Introduction

Acylcarnitines are a class of molecules generated when L-carnitine forms esters (and theoretically amides) with diverse acyl groups, including those derived from fatty acids, amino acids, xenobiotics, and microbial products. Canonically, carnitine functions in the transport of long-chain fatty acids into mitochondria for β-oxidation. Accordingly, fatty acylcarnitines are widely used as indicators of mitochondrial β-oxidation flux and overall fatty acid metabolism, processes central to ATP production and energy homeostasis^1,2^. However, accumulating evidence indicates that acylcarnitines have broader metabolic roles. Beyond fatty acid oxidation, they participate in amino-acid and nitrogen catabolism, peroxisomal and mitochondrial oxidation of diet-derived branched-chain fatty acids, maintenance of CoA and redox homeostasis, endocrine regulation, detoxification, and inter-organ metabolic signaling^3–7^. Distinct acylcarnitine profiles are also clinically important: they are routinely used to diagnose inborn errors of metabolism and have been associated with cardiometabolic, neurological, as well as inflammatory diseases^8–10^. Due to their broadening relevance, carnitine derivatives are increasingly being investigated as biomarkers and therapeutic agents in clinical trials^7,8,10^. Despite their broad physiological and clinical relevance, the full chemical diversity of acylcarnitines remains incompletely characterized in untargeted metabolomic data sets, in part because the range of possible acyl modifications and molecular scaffolds that can conjugate to carnitines is poorly defined.

In recent work, we generated an MS/MS spectral reference library using multiplexed organic synthesis, which included an ibuprofen-carnitine conjugate among the synthesized products^11^. Using a reverse metabolomics approach^12,13^, we subsequently observed this ibuprofen-carnitine species in human LC-MS/MS datasets, revealing that xenobiotic-derived acylcarnitines are already present but largely unrecognized in existing data. In a mouse injury model this ibuprofen-carnitine explains delayed muscle injury recovery^11^. These findings highlight that acylcarnitine biology extends far beyond classical lipid metabolism into environmental and drug metabolism raising a broader question: how many additional acylcarnitines are already present in public untargeted metabolomics datasets but remain unannotated? Given that acylcarnitines ionize efficiently in positive mode and display characteristic fragment ions^14^, we reasoned that those MS/MS fingerprints could be leveraged to mine existing metabolomics repositories for both known and yet-to-be characterized acylcarnitines.

To address this gap, we set out to build a pan-repository acylcarnitine MS/MS spectral library that enables the systematic detection of known and previously uncharacterized molecular species across untargeted metabolomics datasets. The recently developed Mass Spectrometry Query Language (MassQL)^15^ allows systematic filtering of MS/MS data for defined fragmentation patterns. We have previously applied this approach to generate class-specific MS/MS libraries for *N*-acyl lipids, bile acids, and glucuronidated metabolites^16–18^; however, those efforts were restricted to a selected set of Orbitrap-specific datasets within a single repository (GNPS/MassIVE)^19^, or based on specific experimental designs. We have now extended this framework to include MetaboLights^20^ and Metabolomics Workbench^21^, allowing pan-repository MassQL querying and dissemination of results through FASST/MASST and PanReDU^22–25^. Leveraging the diagnostic fragmentation behavior of carnitines, we extracted MS/MS spectra for thousands of candidate acylcarnitines and curated them into an unified open library that more than doubles the number of carnitines documented to date. Coupled with reverse metabolomics^12,13^, this resource enables both structural discovery and biological contextualization of acylcarnitines - thereby expanding the acylcarnitine landscape and providing an enhanced foundation for chemical annotation and functional interpretation of this growing metabolite class.

## Results

### Detection of carnitines in public metabolomics data

Despite the important biological roles of carnitines, no single resource currently compiles the structures of all known and yet-to-be-described carnitines. This makes it difficult to estimate their overall diversity or systematically annotate them in metabolomics studies. To get a sense of how many MS/MS of acylcarnitines might exist, the Human Metabolome Database (HMDB 5.0)^26^ reports 1,199 catalogued carnitines, of which 71 have been experimentally detected and/or quantified in biological samples, while the remainder are *in silico* predictions. PubChem^27^ lists 723 entries containing “carnitine”, encompassing naturally occurring, isotopically labeled standards, assay results with carnitine-related enzymes, and synthetic derivatives. In addition, large-scale metabolomics surveys have further identified hundreds of different carnitine mass spectrometry signatures across tissues, biofluids, and model organisms, reflecting the chemical diversity of this metabolite class. Notably, the two largest carnitine discovery mass spectrometry studies reported the detection of 586 (representing 157 different conjugations) and 1,136 (representing 398 modifications) mass spectrometry signatures of carnitines^14,28^. Finally, a recent study synthesized 76 acyl carnitines of saturated, unsaturated, dicarboxylated and hydroxylated fatty acids^29^ with up to 21 carbons for reverse metabolomics analysis and establishing their use for classification of health status.

Based on these observations, we hypothesized that many additional carnitine derivatives (or carnitine MS/MS signatures) remain unreported in existing metabolomics datasets. Leveraging our recently developed capability to filter MS/MS patterns across metabolomics data repositories, we set out to build an acylcarnitine MS/MS reference spectral library to enable retrospective analyses and support future studies of this important metabolite class. In our previous MassQL-based efforts for bile acids and *N*-acyl lipids,^16,17^ queries could only be executed within the GNPS/MassIVE repository, reflecting the computational and software constraints at that time. Recent hardware scaling and software-engineering upgrades in the GNPS2 infrastructure now allow MS/MS pattern queries to be executed across GNPS/MassIVE, MetaboLights, and the Metabolomics Workbench, thereby substantially expanding the searchable space of public data^11,24^. Initially, we applied MassQL to filter MS/MS spectra potentially originating from carnitines across nearly 2 billion spectra from these repositories that were available in May/June of 2025. All repository-scale searches were restricted to positive-ion mode because acylcarnitines contain a quaternary ammonium group that carries a permanent positive charge, leading to efficient ionization and robust formation of the diagnostic product ions at *m/z* 60.0808, 85.0284, and 144.1019 under electrospray in positive mode^14^ (**Supplementary Figure S1a**). In addition, the majority of LC-MS/MS metabolomics datasets currently deposited in public repositories were acquired in positive ionization mode, so focusing on this polarity both aligns with the intrinsic ionization properties of carnitines and maximizes the practical coverage of our MassQL-based search strategy.

We first validated this query using the standard GNPS spectral libraries^19^. In this analysis, the query retrieved 2.1% spectra of compounds that were not carnitines. Applying the query against the three repositories mentioned above, we retrieved 1,402,842 spectra. Matches to data-independent acquisition (DIA) mode data were removed, and we used the Falcon clustering algorithm to reduce data redundancy^30^, resulting in 37,215 clustered MS/MS spectra. The estimated FDR was 2.4% based on spectral library matching post-clustering, and we used classical molecular networking^31^ to remove all the MS/MS spectra of sub-networks for the nodes that contained one or more non-carnitine annotations. Finally, we retained only carnitine modifications (referred to as delta masses) that were observed in at least two independent studies. The final library comprised 34,222 spectra retrieved from 707 datasets and representing 2,857 unique delta masses, forming the basis of an MS/MS reference collection to aid downstream annotation of candidate carnitine modifications (**Figure 1a**, **Supplementary Figure S1**). To assist in the interpretation of these modifications that carnitine can undergo, we used *in silico* molecular formula predictions, and the ones that obtained modifications solely with C, H, and O atoms we assigned a putative explanation based on LIPID MAPS^32^ standard annotation nomenclature for carnitines (e.g., CAR C18:1;O would be interpreted as a carnitine conjugated with a chain length of 18 carbons, which could be linear or branched, containing one unsaturation and one hydroxylation - **Figure 1b**).

**Figure 1.**
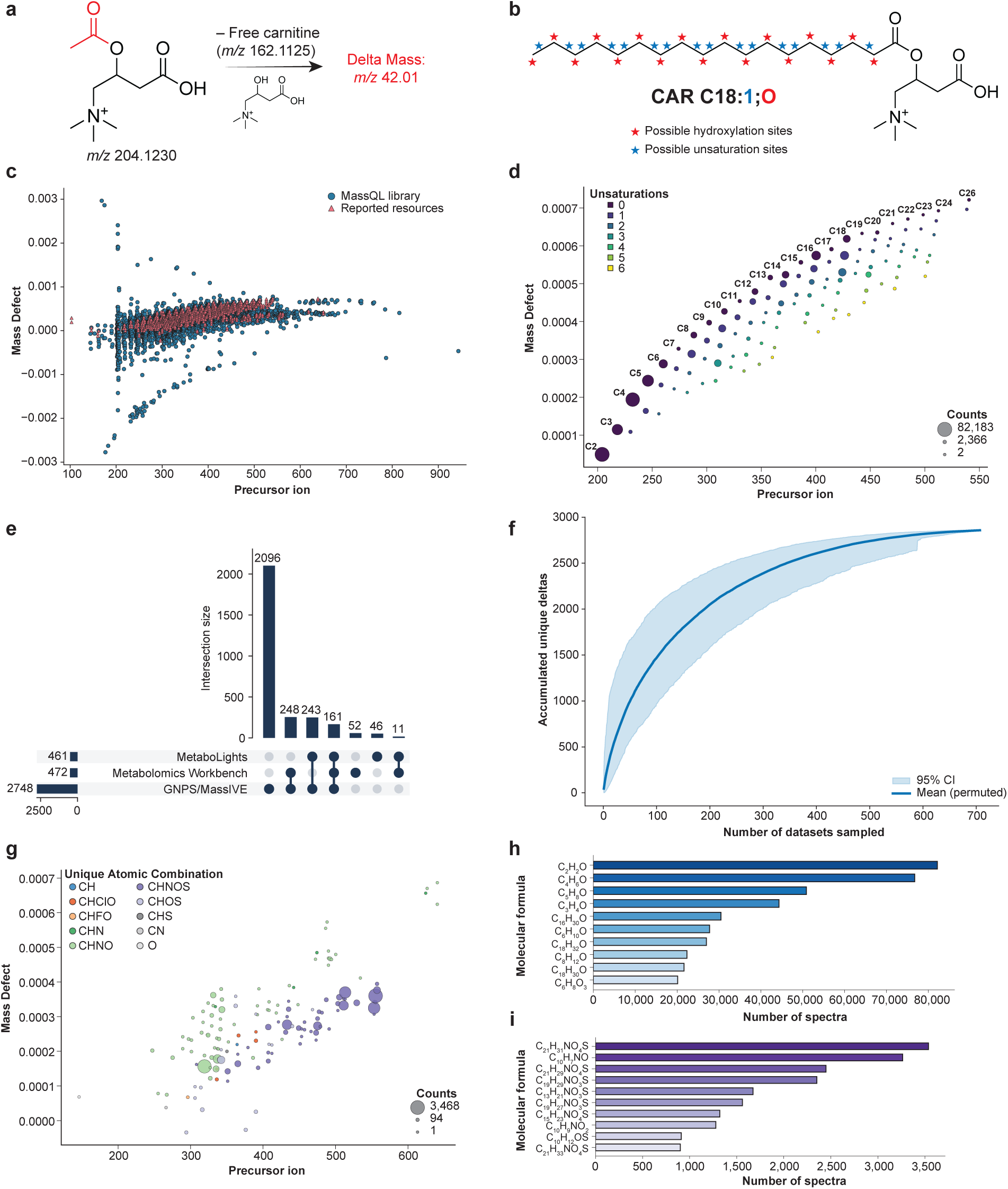
Coverage of the carnitine library. **(a)** Representation of the delta mass calculation using acetylcarnitine as an example. A subtraction of the unconjugated carnitine will result in the delta mass 42.01, which corresponds to an acetylation. **(b)** Representative example of the uncertainty of the position of the hydroxylation and unsaturation for CAR C18:1;O. The 18-carbon chain can be linear or ramified**. (c)** Mass defect plot of the modifications part of the library created in this work, compared with resources currently available, including three large mass spectrometry studies^14,28,29^, LIPID MAPS^32^, PubChem^27^, ChEBI^35^, and HMDB^26^. **(d)** Mass defect plots of the modifications with atomic composition putatively consisting exclusively of C, H, and O atoms. **(e)** UpSet plot of the distribution of the detected carnitine modifications across the metabolomics repositories used in this study. **(f)** Rarefaction curve displaying the number of carnitine analogs against the number of datasets sampled. **(g)** Mass defect plot showing delta masses excluding modifications containing only C, H, and O. **(h)** Frequency of the most common elemental compositions among all modifications. **(i)** Frequency of elemental compositions after removing CHO-only modifications.

In principle, the MS/MS spectra that constitute the library could represent in-source fragments of the acyl substituents, but not of the carnitine moiety itself, as fragmentation of carnitine would disrupt its characteristic diagnostic fragment ions. These signals could also arise from alternative ion species, such as adducts or multimers. However, this scenario is also less likely because carnitines typically do not form stable Na⁺, K⁺, or similar adducts under standard electrospray conditions. If such species were to form, and we are not eliminating this possibility without further evidence, they would likely involve multiply charged ions, which are not represented in available reference MS/MS libraries and may be overlooked. Unlike many metabolites that rely on adduct formation for efficient ionization, carnitines contain a permanent quaternary ammonium group and therefore readily ionize as intact molecular ions. Another possibility is that acylation occurs via electrospray droplet chemistry^33^, which would imply post-chromatographic conjugation reactions. This could only occur if free carnitine co-migrates chromatographically with the acyl donors (e.g., fatty acids). Such a scenario is improbable for most of the MS/MS spectra we observed, particularly given the diversity and generally less polar nature of the acyl groups that would not co-migrate with carnitine. It could, however, play a role if an acylating molecule has the same polarity and chromatographic behavior as carnitine. Instead, many carnitine MS/MS spectra exhibit systematic patterns in *m/z* and mass defect that form homologous series consistent with defined chemical substitutions. These include variations in acyl chain length, degree of unsaturation, hydroxylation, and, within the same dataset, varying retention times. These patterns align well with known features of fatty acid metabolism, including β-oxidation and odd-chain fatty acid pathways^34^.

Mass defect plots showed that our library offers a major expansion of the known chemical space of carnitines when compared to other resources currently available^14,28,29^ (**Figure 1c**). We observed that even chain substitutions are more frequently detected than the odd chain ones, regardless of the hydroxylation state (**Figure 1d, Supplementary Figure S2**), and that most of the unique delta masses were retrieved from the GNPS/MassIVE repository (**Figure 1e**). This is consistent with GNPS/MassIVE containing the largest number of MS/MS spectra among the three repositories^24^. To estimate the potential of discovery of new carnitine modifications, we created a rarefaction curve, which shows a rapid increase in accumulation of new unique delta masses with diminishing returns as more datasets are sampled, reaching close to a plateau when we reach 700 sampled datasets (**Figure 1f**). When datasets using alternative extraction methods or chromatography conditions are incorporated, we may observe a further increase; however, under the current broad range of experimental conditions, these results suggest a diminishing likelihood of acquiring data on new carnitines that have unique atomic compositions (though they may still be structurally unique) as additional samples are added. While a significant portion of our library contains atom combinations such as CHNO and CHOS (**Figure 1g,i**), most of the putative modifications retrieved contain only the atoms CHO within a range of 2 to 26 carbons and up to 6 unsaturations (**Figure 1d,h**). The most frequent modifications matched to public data were conjugations corresponding to C2 and C4 short-chain fatty acids, observed 82,183 and 76,781 times, respectively.

To implement a reverse metabolomics strategy and systematically link acylcarnitine MS/MS spectra to tissues, organisms, and study contexts, we performed repository-scale searches using fastMASST (FASST)^22,23^, an accelerated implementation of MASST, across GNPS/MassIVE, MetaboLights, and Metabolomics Workbench. Because our query included a fragment at low *m/z* (60.0808) - which often falls outside the MS/MS acquisition range of many datasets - using MASST’s cosine-similarity matching (which tolerates low-abundance and missing ions) increased our ability to recover carnitine matches present in additional datasets. Using this approach, we obtained matches to 3,872,050 MS/MS spectra from 220,793 files across 1,977 datasets, underscoring the widespread detection of carnitines in public metabolomics repositories. Among datasets with curated, standardized controlled-vocabulary metadata in PanReDU^24,25^, we mapped 2,329 carnitine modifications (delta mass values) in humans and 1,739 in rodents (**Figure 2a,b**), revealing a broad distribution of candidate carnitine analogs with high prevalence in urine, feces, blood, and gastrointestinal tract samples. Additional tissues showing several potential carnitine analogs included heart, placenta, and lung in rodents, and milk and the alveolar system in humans. Furthermore, unique delta masses were also tissue/biofluid-specific, as summarized by the UpSet plots (**Figure 2c,d**). In human samples, 423 delta masses were observed exclusively in urine, followed by sets shared between urine and blood, then those unique to feces, and finally those shared among all three (**Figure 2c**). In rodents, the limited availability of urine samples yielded only 59 unique delta masses, with feces exhibiting the highest diversity of carnitines (**Figure 2d**). Because rodent datasets include a wider range of tissues, they also provide a more complete map of body-part distribution than is currently available for humans. In both, we also observed many carnitines that are shared across multiple tissues and biofluids.

**Figure 2.**
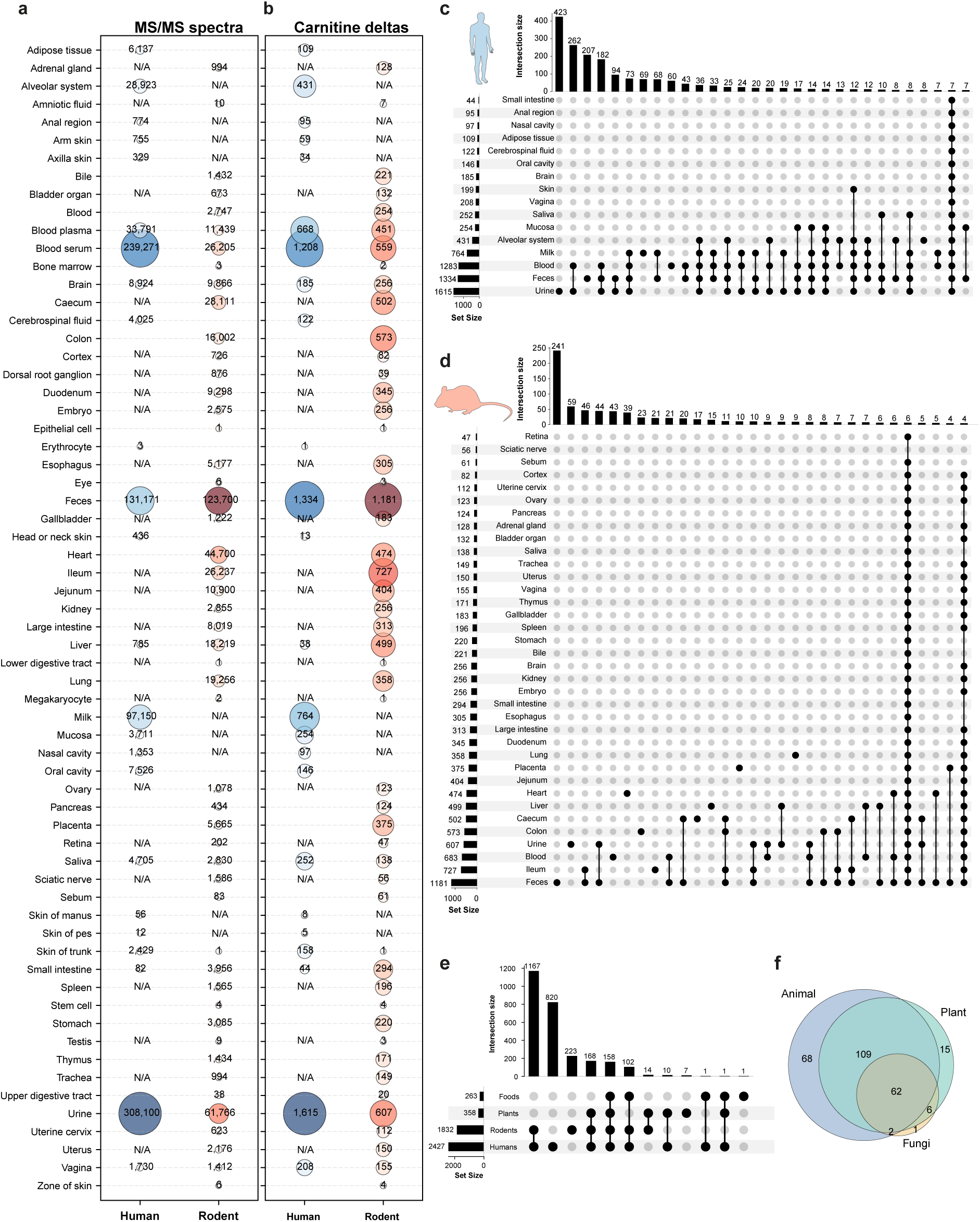
Repository distribution of candidate carnitines, represented as delta masses, from FASST searches. **(a,b)** Repository coverage showing **(a)** MS/MS spectra counts and **(b)** unique carnitine delta masses across human and rodent tissues or biofluids. Numbers in blue and red circles indicate measurements for humans and rodents, respectively; N/A indicates no data available in the repository. **(c,d)** UpSet plots illustrating tissue/biofluid-specific and shared distribution patterns of carnitine delta masses in **(c)** humans and **(d)** rodents. For clarity, only the top 30 intersections are shown. Horizontal bars indicate the number of delta masses detected in each tissue/biofluid; vertical bars show the number of delta masses shared across tissue combinations, with filled circles below indicating which tissues are included in each intersection. **(e)** Cross-kingdom distribution of carnitines across humans, rodents, plants, and food samples. We did not show matches to microbial data, as growth media often contain carnitines, and although microbial metabolites such as short-chain fatty acids, or microbial metabolites of phenylalanine are known to be conjugated to carnitine^38^, it was not possible to evaluate if the conjugation was carried out by microbes or not. **(f)** Number of modifications observed for each food type at a level 1 of food ontology according to foodMASST^39^. Icons were obtained from Bioicons.com

Out of the 2,857 atomic compositions we found, 358 and 263 were detected in samples from plants and foods, respectively (**Figure 2e**). Although carnitines are well known to be produced by mammalian systems, they can also be consumed as part of the diet (primarily foods of animal origin, **Figure 2f**), and are also produced by plants^36^. To test whether microbes further influence carnitine levels, we re-analyzed published datasets involving microbiome perturbations using our library. In a mouse study in which animals received a cocktail of antibiotics, all of the carnitine derivatives matches with our library showed increased levels as microbes are depleted, consistent with these molecules being metabolized by the microbiome or that the microbiome facilitates their absorption (**Supplementary Figure S3a**). In line with this, prior studies have reported increased levels of a variety of acylcarnitines in maternal feces following antepartum or postpartum ampicillin exposure in mice^37^. We then examined a second dataset comparing germ-free mice to specific-pathogen-free (SPF) mice, mice colonized with segmented filamentous bacteria (SFB), and mice monocolonized with individual gut microbes. Carnitine derivatives showed group-specific alterations relative to germ-free controls, indicating that the microbiome can selectively shape the carnitine pool (**Supplementary Figure S3b**). Additionally, we observed a general trend of increasing abundance from C2 to C16 species upon colonization, which was more pronounced in colon samples than in the small intestine.

### Utility of the carnitines library

To further highlight the utility of the carnitine MS/MS library, we illustrate three applications. In the first, we applied the library to a time-restricted feeding study in mice, spanning 20 h and covering 14 organs, gastrointestinal (GI) contents, and biofluids^40^. We obtained 162 matches with our library that passed both a cosine similarity match and a MassQL filter^15,41^ for carnitine diagnostic ions. Of these, 131 had molecular formula annotations relative to CHO modifications, all of which also had within-sample different retention times, consistent with pre-chromatography rather than post-chromatography formation of the acylcarnitines, which were considered for downstream analyses. This analysis revealed patterns in carnitine metabolism, underscoring the roles of the diurnal and feeding cycles in systemic energy homeostasis. Along the GI tract, peak times shifted as contents progressed: long-chain conjugates peaked at 8 h from the stomach to the ileum, while cecal and colon contents showed peaks at 4 and 12 h (**Figure 3a**). Blood, heart, and skin were enriched in short-chain conjugates; the eye and brain showed minimal temporal variation. Urine was the most distinct matrix, with higher levels of medium-chain conjugates peaking at 8 h. It also showed the most different structural profiles, with enrichment of odd-chain species, one to two unsaturations, and two hydroxylations (**Supplementary Figure S4**), whereas most other tissues and biofluids were dominated by even carbon chains, without unsaturations or hydroxylations. We further identified synchronized groups of carnitine analogs (**Figure 3b,c**), including a urine-specific cluster of 64 compounds peaking at 8 h. Multiple tissues contained clusters peaking at different times. For example, duodenal contents showed one cluster peaking at 4 h and another at 16 h, while the kidney demonstrated clusters peaking at 8 h and 20 h. The largest synchronized carnitine cluster in blood reached maximum levels at 12 h.

**Figure 3.**
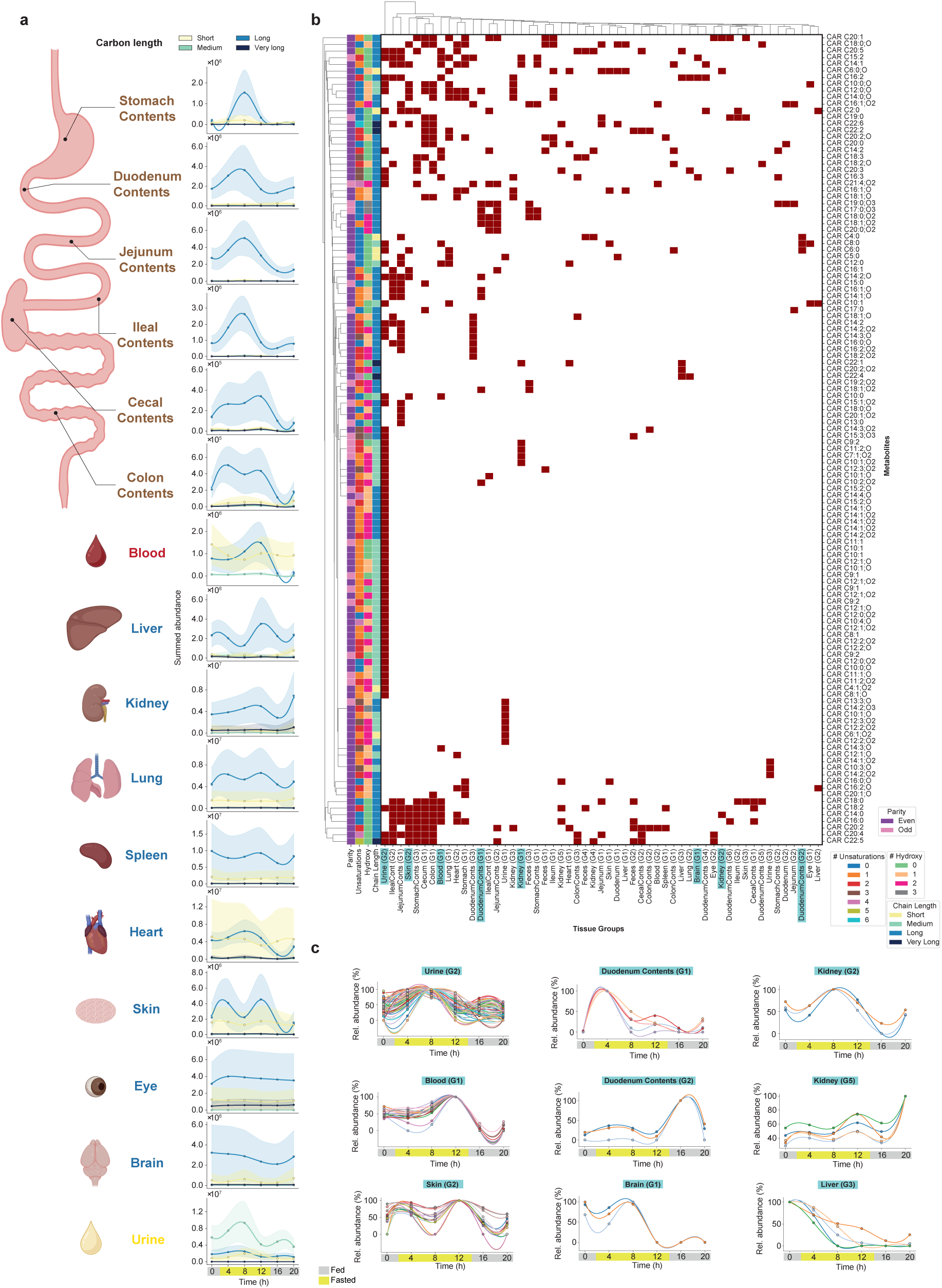
Diurnal patterns of carnitine derivatives across multiple tissues and biofluids in time-restricted feeding mice. **(a)** Temporal profiles of total carnitine abundance grouped by carbon chain length of the substitution (short: C2-C6; medium: C7-C12; long: C13-C21; very long: C22-C30) across 14 tissue types, gut contents, and biofluids sampled at six time points over a 20-hour period. Solid lines represent smoothed cubic spline fits of summed abundance values with Gaussian filtering for each chain length group. Shaded bands indicate 95% confidence intervals calculated from the standard error of the sum. Individual points show the sum of all detected carnitines within each chain length category at each time point. **(b)** Heatmap showing hierarchical clustering of individual carnitine modifications (rows) across tissue-specific temporal groups (columns). Dark red squares indicate the presence of specific carnitines within each cluster. Left annotation bars denote structural information (parity, number of unsaturations, number of hydroxyl groups, and chain length group). Tissue groups (G1, G2, etc.) represent distinct temporal clusters identified within each tissue, demonstrating coordinated regulation of carnitine subsets. **(c)** Representative temporal dynamics of selected tissue-specific carnitine clusters showing the relative abundance (%) over the 20-hour cycle. Each panel displays multiple carnitine species (colored lines) within a given cluster, illustrating synchronized patterns. Icons were obtained from BioRender.

In the second use case, we used the newly created library to annotate carnitines in a previously analyzed untargeted metabolomics dataset, which included longitudinal samples from 55 mother-infant dyads from Bangladesh^42^ (MSV000096943). Briefly, the dataset included infant plasma (n = 34) and stool (n = 379) samples, and maternal milk samples (n = 119) collected at multiple timepoints from birth to around six months postpartum. A total of 300 features were annotated as putative carnitines, of which 176 had putative CHO modifications. Infant fecal samples had the most enriched carnitine pool, followed by plasma and milk (**Figure 4a**). Many of the annotated carnitines were shared across the three biofluids and spanned a wide range of carbon lengths, hydroxyl groups, and unsaturations (**Figure 4a**). We also investigated the trajectories of acylcarnitines grouped by carbon length, hydroxyl groups, unsaturations, and parity of the carbon numbers within the first 6 months after birth, expanding on what has been shown previously^42^. Minor changes were observed for human milk (**Figure 4b**), while clearer shifts were observed in infant feces (**Figure 4c**). Specifically, we observed a decrease in short-chain acylcarnitines (LME: β = −0.007, p = 0.01), in line with previous literature^43^, alongside an increase in long- (LME: β = 0.009, p = 1.0e-05), or very long-chain (LME: β = 0.026, p = <2e-16), acylcarnitines, as well as acylcarnitines with a higher degree of unsaturation.

**Figure 4.**
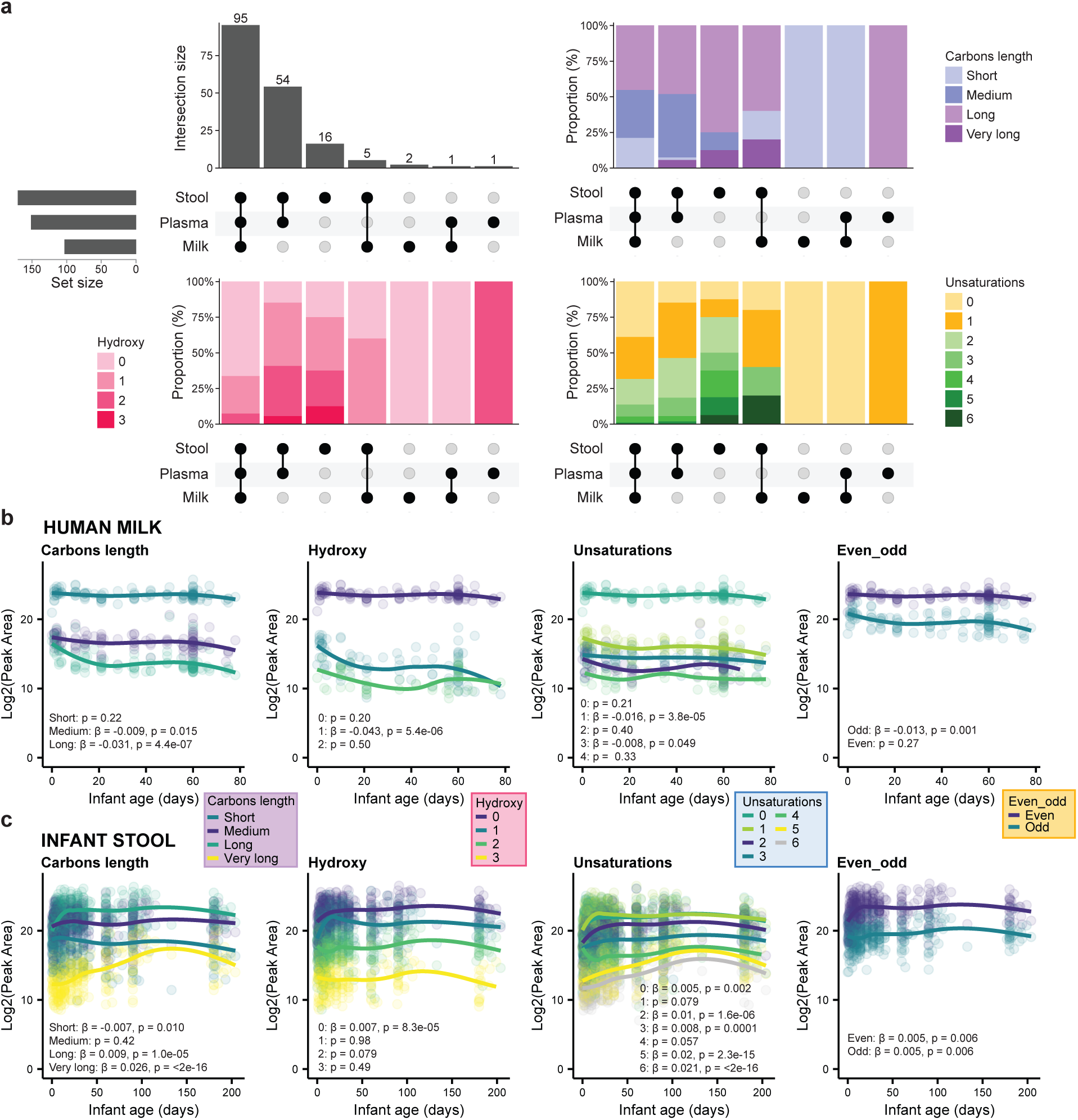
Acylcarnitines in maternal milk and infant plasma/feces: composition and time courses over the first 6 months. **(a)** UpSet plot of annotated acylcarnitines (n = 176) across infant fecal (n = 379) and plasma (n = 34) samples, and human milk (n = 119) samples. The upper left panel shows acylcarnitines unique to or shared between biofluids. The upper right panel shows the distribution by carbon length, the lower left panel by hydroxy groups, and the lower right panel by unsaturations. **(b)** Trajectories for acylcarnitines in human milk by carbon length, hydroxy groups, unsaturations, and parity in the number of carbons, respectively. **(c)** Trajectories for acylcarnitines in infant feces by carbon length, hydroxyl groups, unsaturations, and even or odd number of carbons, respectively. β and p values in **b** and **c** are derived from linear mixed-effect models with subject as a random effect. Zero values were removed from the visualization. Trend lines were added to illustrate patterns.

Given that urine represents one of the richest biofluids for carnitine detection (**Figure 2a,b**; **Figure 3**), we investigated which datasets containing urine samples yielded the highest number of FASST spectral matches **(Figure 5a**). This analysis highlighted the PlusRise Urobiome study (MSV000096359), a cohort of 622 urine samples collected from adult women as part of an investigation of bladder health^44,45^. Applying our spectral library with post-processing MassQL filtering^15,41^ resulted in 413 candidate carnitine spectral matches. Molecular networking and annotations based on spectral similarity revealed three distinct annotation patterns: annotated exclusively by existing public libraries (relative to other classes of molecules), nodes annotated by both our library and existing public libraries, and nodes exclusively annotated as belonging to carnitines by our carnitine library (**Figure 5b,c**). A molecular network with the largest number of annotations obtained with publicly available libraries of known carnitines were also assigned to be carnitines by the acylcarnitine library that we generated in this work. Moreover, we also observed nodes within this same network that matched exclusively our library of carnitine predictions (**Figure 5b**). These are primarily classical fatty acid conjugates of carnitines. There are also networks in which we exclusively observed annotations with our library (**Figure 5c**), that did not fragment similarly to the fatty acid conjugates. These could either be misassigned as MS/MS belonging to the carnitine family, or they could serve as a source of yet-to-be-identified acylcarnitines. Reverse cosine applied to one of such networks (**Figure 5c**) suggested that conjugation of carnitine with phenylpropanoid-like groups (**Supplementary Table S1**), and if this prediction is correct, these would represent a class of acylcarnitines not previously reported^11,45,46^. Mass differences between nodes were consistent with common phenylpropanoid-type modifications, including degrees of unsaturation (Δ2.015 Da), hydroxylation (Δ15.995 Da), methylation (Δ14.016 Da), and methoxylation (Δ30.011 Da). Notably, we also observed mass differences consistent with glucuronidation (Δ176.032 Da) and sulfation (Δ79.957 Da).

**Figure 5.**
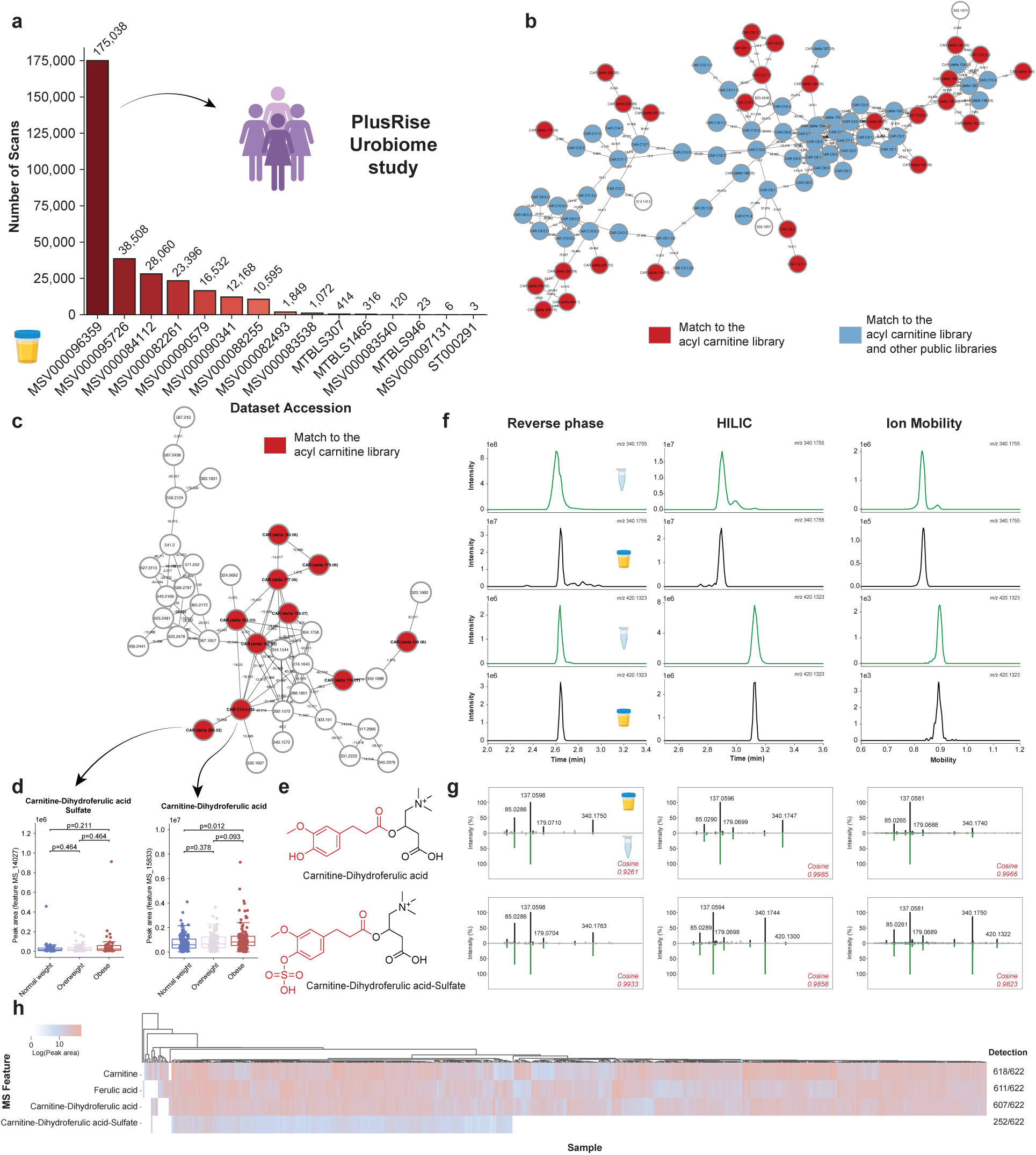
Phenylpropanoid-carnitine conjugates detected and validated in human urine. **(a)** Public datasets containing urine samples ranked by the number of FASST matches to the acylcarnitine spectral library, highlighting the public PlusRise Urobiome study^44^ selected for downstream analysis. **(b)** Feature-based molecular network^49^ of carnitine-related metabolites detected in this dataset. Nodes represent MS/MS features and edges represent spectral similarity; nodes are colored by annotation source: acylcarnitine library only (red) or shared with existing public libraries (blue). **(c)** Subnetwork annotated exclusively by the acylcarnitine library, suggesting candidate phenylpropanoid-derived acylcarnitines based on reverse cosine^45,46^ relationships and characteristic mass differences. **(d)** Relative abundance of carnitine-dihydroferulic acid and carnitine-dihydroferulic acid sulfate across BMI groups (normal weight, n = 142; overweight, n = 110; obese, n = 136). Zero values were removed from the analysis. Significance was tested using the non-parametric Mann-Whitney U test and p values were corrected for multiple comparisons using the Benjamini-Hochberg correction. Boxplots indicate the first (lower), median, and third (upper) quartiles, while whiskers are 1.5 times the interquartile range. **(e)** Structures of synthesized standards used for validation. **(f)** Orthogonal validation showing co-elution under reverse-phase LC, separation by HILIC, and matching ion mobility drift times between standards and biological features. **(g)** MS/MS spectral comparisons between synthetic standards and biological features. **(h)** Heatmap showing detection patterns of the validated compounds across the cohort. Icons were obtained from BioRender.

To provide additional support for some of these annotation predictions, we selected two candidate structures for chemical synthesis: a carnitine-dihydroferulic acid conjugate and its sulfated analog (**Figure 5d,e**). The sulfated species was prioritized because sulfated acylcarnitines are poorly documented despite sulfated metabolites being commonly detected in urine. Notably, carnitine-dihydroferulic acid levels were elevated in participants with obesity compared to normal weight individuals, while its sulfated analog showed a similar trend that did not reach statistical significance (**Figure 5d**). Initial chromatographic analysis under reverse-phase conditions revealed that both compounds co-eluted, and retention time separation could not be achieved despite attempts at optimization of chromatographic conditions (**Figure 5f**, left). This observation was further complicated by the modification being a sulfate. Sulfate groups are labile under electrospray ionization and do not require much energy input to break the sulfodiester bond. This raised the possibility that the non-sulfated feature detected in the biological sample could be an in-source fragment of the sulfated compound rather than a distinct compound. To resolve this ambiguity, we switched to hydrophilic interaction chromatography (HILIC), which achieved baseline separation of the two compounds and enabled retention time matching of the two compounds against the biological sample (**Figure 5f**, center). Ion mobility further provided orthogonal confirmation, with both synthetic standards and biological features resulting in matched ion mobility drift times (**Figure 5f**, right). In all conditions, MS/MS spectral matching produced high cosine similarity scores (**Figure 5g**). The combined results give rise to level 1 annotation according to the Metabolomics Standards Initiative^47^ for both the sulfated and non sulfated dihydroferulic acid carnitine. Finally, examination of feature prevalence across the PlusRise cohort revealed that carnitine-dihydroferulic acid was detected in 607 participants, and carnitine-dihydroferulic acid-sulfate in 252 participants (**Figure 5h**), indicating that these previously unreported conjugates can be widespread in biological studies. The broad detection of ferulic acid and its isomers, which are primarily food-derived, is consistent with their widespread occurrence in foods when analyzed using StructureMASST^48^. Using the ferulic acid structure as input against FASSTrecords yielded matches in public datasets associated with foods, confirming its presence across a broad range of dietary sources - such as legumes; grains including quinoa, corn, wheat, and rice; citrus fruits; vegetables such as broccoli, cabbage, brussels sprouts, beet, potato, sweet potato, garlic, and onion; as well as tea and a variety of herbs and spices (**Supplementary Figure S5**). Given that these are all commonly consumed foods, the widespread detection of ferulic acid-derived metabolites is not unexpected. The novel finding in this study, however, is that these metabolites are observed as carnitine conjugates, a feature that has not previously been reported.

## Discussion

In this study, we expand the detectable MS/MS landscape of acylcarnitines, revealing a level of structural and contextual diversity that is represented by the MS/MS that is not captured by conventional targeted or untargeted metabolomics workflows. By learning from reference MS/MS spectra and systematic interrogation of public metabolomics repositories, we show that acylcarnitines extend beyond their canonical roles in fatty acid transport and energy metabolism. Instead, they function as a broadly deployable conjugation framework linking endogenous metabolism with diet, microbiome derived metabolism, and xenobiotic exposure.

Relative to existing carnitine-focused resources^14,28,29^, our approach shifts from narrowly sampled reference materials or synthetic approaches toward a pan-repository view of the acylcarnitine chemical space. Previous work demonstrated that MS/MS pattern-based filtering using MassQL can expand annotation for specific metabolite classes within individual repositories^16,17^. Here, we generalize this strategy across GNPS/MassIVE, MetaboLights and Metabolomics Workbench, encompassing a broader diversity of instruments, analytical conditions and biological systems. This expanded scope not only increases the number of distinct MS/MS that belong to acylcarnitines, but also captures context-specific variants that are largely absent from conventional reference libraries.

To place this expanded chemical space into biological context, we combined FASST/MASST spectral searches with standardized ReDU metadata. Searching the acylcarnitine library against pre-indexed repository spectra yielded over 3.8 million matches across nearly 2,000 datasets. Mapping these matches to harmonized metadata enables association with organismal origin, sample type and experimental context, effectively transforming the library from a static catalog into a context-aware resource. This integration allows researchers to ask not only which acylcarnitines are present, but where they occur and under which physiological, dietary or environmental conditions, facilitating hypothesis generation and prioritization for targeted validation.

We demonstrate the practical utility of this resource across three biologically distinct applications. In a murine time-restricted feeding study, the library revealed tissue-specific partitioning and temporal dynamics of acylcarnitines across a feeding-fasting cycle. Re-analysis of a longitudinal mother-infant cohort showed that the same resource can be applied retrospectively to interrogate early-life acylcarnitine metabolism across plasma, stool and human milk. Finally, a urine cohort of approximately 600 samples from adult women, revealed molecular networks of carnitine-related features not represented in existing public spectral libraries, where food derived metabolites are conjugated. Together, these examples illustrate that the library can be broadly overlaid onto animal experiments, human cohorts and archived metabolomics datasets.

As with any MS/MS-based resource, appropriate interpretation requires recognition of its strengths and limitations. Spectral matching enables systematic detection of modified and previously unreported acylcarnitines and often distinguishes among classes of acyl modifications, such as differences in chain length, saturation or oxidation state. However, positional and stereochemical isomers require additional orthogonal information, including retention-time matching and/or ion-mobility measurements, for definitive structural assignment. While molecular formulas could be assigned to most library spectra using current prediction tools^50,51^, certain heteroatom-rich modifications remain challenging, underscoring the importance of manual inspection of diagnostic fragments. Applied with these considerations, the library reveals previously unrecognized dimensions of carnitine biology. Importantly, both the MassQL query and the spectral library are reusable, enabling analyses from single experiments to repository-scale studies.

More broadly, this acylcarnitine resource, together with related efforts for bile acids and *N*-acyl lipids^16,17^, illustrates the cumulative value of open, repository-scale metabolomics. A substantial fraction of MS/MS spectra in untargeted studies remains unannotated not because these molecules are inherently obscure, but because existing libraries incompletely sample chemical space - referred to as the dark matter of the metabolome^52,53^. Systematic mining of public datasets for recurring fragmentation motifs and their translation into reusable resources provides a scalable path toward converting this dark fraction into structured, interpretable knowledge. These findings underscore the importance of open data sharing and standardized metadata, as each deposited dataset expands the collective capacity for metabolite annotation and biological insight.

## Summary/Conclusion

In summary, the breadth of carnitine biology only becomes apparent through repository-scale analysis of untargeted metabolomics data. By integrating MS/MS pattern mining with harmonized metadata across public resources, we reveal carnitines as a pervasive and flexible metabolic conjugation system spanning diverse biological contexts. This work highlights repository-scale metabolomics as a powerful approach for discovering and interpreting metabolic diversity that is otherwise dispersed across individual studies.

## Limitations of the study

Despite its breadth, the carnitine MS/MS resource described here has important limitations. Our MassQL queries rely on canonical known diagnostic fragments of the carnitine core and intensity thresholds. This means that carnitines whose fragmentation suppresses these ions or redistributes intensity across alternative fragments may not be captured, even when present in the data. As illustrated in **Figure 5c** (white nodes), a dataset uploaded after generation of the current MassQL library contains multiple MS/MS spectra that match against the MassQL carnitine MS/MS library, yet many others in that subcluster were not captured by the existing query. Lowering the *m/z* 85.0284 intensity threshold from 50% to 5% in the MassQL query allowed nearly all other ions in that network to be recognized as carnitine signatures, indicating that mismatches occur more frequently when the attached acyl side chain undergoes internal fragmentation (**Supplementary Figure S6**).

A comparable pattern is observed in a recently reported set of 81 spectra from 76 chemically synthesized acylcarnitines^29^. In that library, twelve spectra, corresponding to 3-hydroxylated acylcarnitines, received no annotation and failed post molecular networking MassQL filtering^41^ even after relaxing the *m/z* 85.0284 threshold to 5%. Manual inspection revealed a consistent fragmentation pattern: the *m/z* 144.1019 fragment, that is traditionally considered a diagnostic carnitine fragment ion^14^, is absent, the *m/z* 85.0284 ion is barely detectable, and the dominant fragment on those carnitines arises from internal fragmentation at the hydroxylated fatty-acid side chain. Similar internal fragmentation is likely to occur with other conjugates, reducing the intensity of canonical ions and limiting the spectra that is captured by the query.

Therefore, while this work introduces a carnitine detection resource over ten times larger than any previous efforts, substantially expanding the annotatable acylcarnitine landscape, these missed annotations also reveal a clear future opportunity for the field in the discovery of additional acylcarnitines not only by further assigning the structures of observed acyl substituents but also discovering additional ones. Future efforts could expand query sets, develop subclass-specific fragmentation rules, and leverage approaches such as targeted synthesis and machine learning to identify a broader range of acylcarnitine MS/MS spectra. Advancing these strategies will enable the community to refine reference libraries and improve the annotation of this important metabolite class, while also expanding our understanding of the role of acylcarnitines in health and disease.

## Authors contributions

H.M-R., P.C.F, M.v.F., and P.C.D designed the query and performed repository-scale searches.

H.M-R., P.C.F, and W.D.G.N. created the library.

H.M-R., P.C.F, K.E.K, and H.G. performed data analysis.

A.P. performed the synthesis reactions.

P.C.D. and D.S. supervised the synthesis.

L.A.B. acquired urine samples from PLUS Consortium.

H.M-R, A.P., V.D., J.A., and J.Z. acquired LC-MS/MS data.

S.X. performed reverse cosine searches.

E.R.R, I.K., and A.D.P. supervised and acquired the time-restricted feeding study.

H.M-R, P.C.F, K.E.K, and P.C.D wrote the manuscript.

P.C.D. acquired funding and supervised the project.

All authors reviewed the manuscript.

## Acknowledgements

We thank the support from NIH (NIDDK) for the Collaborative Microbial Metabolite Center (U24DK133658) and for supporting the development of tools for structure elucidation (R01DK136117). ERR was supported by PSU/NIDDK (T32DK120509). P.C.P acknowledges the Chen Zuckerberg Initiative (CZI) for the financial support. P.C.F. acknowledges the São Paulo Research Foundation (FAPESP) (grant 2024/17170-0) for funding. ADP was supported by NIH grants (R35ES035027), the PA Department of Health using Tobacco CURE funds, and the USDA National Institute of Food and Federal Appropriations under Project PEN04917 Accession 7006412. SMT acknowledges funding from the Eunice Kennedy Shriver National Institute of Child Health and Development (1P50HD106463).

## Methods

### Design of the MassQL query and creation of the acylcarnitines library

Spectra containing MS/MS diagnostic ions for carnitines were searched among the three main metabolomics repositories: GNPS/MassIVE (which includes datasets grouped by instrument type: QE, Orbitrap, and QToF), MetaboLights, and Metabolomics Workbench. These searches were conducted using the Mass Spectrometry Query Language (MassQL)^15^ across approximately 2 billion publicly available MS/MS spectra (**Supplementary Figure S1a**). Given the widespread availability of datasets acquired in positive ionization mode - and the fact that carnitines ionize more efficiently under these conditions - the query was designed for positive ionization mode.

*QUERY scaninfo(MS2DATA) WHERE MS2PROD=60.0808:TOLERANCEPPM=20:INTENSITYPERCENT=1 AND MS2PROD=85.0284:TOLERANCEPPM=20:INTENSITYPERCENT=50 AND MS2PROD=144.1019:TOLERANCEPPM=20:INTENSITYPERCENT=1*

The false discovery rate (FDR) of the designed query was estimated by running it against the GNPS public spectral libraries, which contain 587,917 spectra representing a wide range of compound classes. The results, available at https://proteomics2.ucsd.edu/ProteoSAFe/status.jsp?task=fc9c9807f0164565b0fcd9c2cf0ac93, resulted in an estimated FDR of 2.12%.

During analysis, we observed that several datasets in the MassIVE QToF collection exhibited the diagnostic ion at m/z 85.0284 at lower relative abundances. To account for this, the intensity threshold for this ion was reduced to 10% for the QToF-specific queries. All queries were executed using the GNPS2 MassQL workflow, and the results can be accessed through the following links:

- Metabolights: https://gnps2.org/status?task=a61b20c5251242afb60ff193146bd1d0
- Metabolomics Workbench: https://gnps2.org/status?task=268a473d58644fddaafdcd6f03ad94cd
- GNPS/MassIVE (Orbitrap collection): https://gnps2.org/status?task=25a8a3ff61e94fcaad99faa6a556faa5
- GNPS/MassIVE (QE collection): https://gnps2.org/status?task=e9013a4b1e94438f97e501122239e71c
- GNPS/MassIVE (QToF collection): https://gnps2.org/status?task=b27657fb7c854fdd9106bcd6ea446890

An additional observation arose from the inspection of MetaboLights results: an unusually high number of scans was retrieved from dataset MTBLS2295, each containing a large number of fragment ions. Upon closer examination, we found that this dataset was acquired using data-independent acquisition (DIA), and the MS/MS spectra were overall very noisy. Due to these characteristics, spectra from MTBLS2295 were excluded from further analysis. After this filtering step, the resulting MGF file contained a total of 1,317,126 spectra (**Supplementary Figure S1b**). To reduce redundancy among the retrieved MS/MS spectra, we applied the Falcon (version 1.3)^30^ clustering algorithm using the following parameters: precursor ion tolerance of 20 ppm, fragment ion tolerance of 0.02 Da, minimum cosine similarity score (eps) of 0.1, a minimum of 3 fragment peaks per spectrum, a minimum *m/z* of 50, and presence of spectra in at least two samples. The most representative spectrum from each cluster was selected for downstream analysis. Falcon clustering yielded 37,215 unique MS/MS spectra (**Supplementary Figure S1c**). Using this clustered MGF file, we performed a classical molecular networking job (https://gnps2.org/status?task=e765ec00637b4ddbac71d58fca8cc654) for both FDR estimation and network generation. The estimated FDR was 2.37%. From the resulting molecular networks, we removed all spectra associated with nodes that were connected to compounds not annotated as carnitines (**Supplementary Figure S1d**).

Two additional filters were applied at this stage to further refine the library. First, mass defect plots of the delta masses revealed an abnormal cluster of values corresponding to precursor ions between ∼600 and 630 Da, observed exclusively in MetaboLights datasets (**Supplementary Figure S1d**). A closer inspection showed that two datasets (MTBLS2274 and MTBLS1040) were not acquired using data-dependent acquisition, and were therefore excluded. This highlights the importance of consistent data acquisition metadata. Second, we applied a filter to retain only delta masses that appeared in at least two independent datasets (**Supplementary Figure S1d**). While this may exclude modifications that are very unique to a specific study, it increases confidence by focusing on features observed across more than one study. After applying these filters, the final dataset consisted of 34,222 MS/MS spectra, representing 2,857 potential carnitine modifications, which were used to construct the carnitine spectral library (**Supplementary Figure S1e**).

### Chemical space of the library

To compare the modification space captured by the acylcarnitine library with previously reported acylcarnitines (**Figure 1c**), we assembled a reference set of acylcarnitine structures from three MS-based studies^14,28,29^, LIPID MAPS^32^, PubChem^27^, ChEBI^35^, and HMDB^26^. PubChem entries were retrieved by searching for the term “carnitine” in the compound name and subsequently removing records annotated as reagents, salts, or isotopically labelled analogues. From HMDB, we retained only acylcarnitines classified as “detected” or “quantified” in biological samples. For all resources, the mass of the acyl moiety was computed by subtracting the mass of the free carnitine core, and delta masses (Δm) were calculated by rounding to two decimal places and retaining unique values.

For the MassQL-derived acylcarnitine library, Δm for each entry was defined as the difference between the precursor ion mass and the mass of the carnitine core; for literature-derived acylcarnitines, Δm was computed analogously using the molecular mass. The Δm mass defect was calculated as the difference between Δm and its nearest integer mass and plotted against precursor ion mass (for acylcarnitines in the MassQL library) or molecular mass (for literature entries) to generate the mass-defect plots in **Figure 1c**. For **Figure 1d**, we restricted the analysis to library entries with a putative molecular formula explanation in the CAR Cx:y;Oz format and selected only those with O = 0 in the acyl moiety (i.e. putative explanations without oxygen). An analogous plot for oxygen-containing putative explanations is provided as **Supplementary Figure S2**.

To examine how acylcarnitine modifications are distributed across repositories (**Figure 1e**), we created a presence/absence matrix summarizing whether each unique Δm was observed in GNPS/MassIVE, MetaboLights, and/or Metabolomics Workbench. This matrix was visualized as an UpSet plot, summarizing shared and repository-specific Δm values. For the rarefaction analysis (**Figure 1f**), we considered all datasets containing at least one unique Δm and performed 1,000 random permutations of dataset order. For each permutation, we computed the cumulative number of unique Δm values as datasets were added sequentially, and then the mean and 95% confidence interval across permutations were plotted as a function of the number of datasets sampled.

For **Figure 1g**, we focused on library entries that lacked a putative explanation but for which the elemental composition of Δm could be predicted using SIRIUS and/or BUDDY. Each Δm was assigned to a “unique atomic composition” category based on its elemental formula (for instance, CHNO, CHOS, CHNOS), and the Δm defect was plotted against precursor mass. The bar plots in **Figure 1h,i** summarize the frequency of Δm molecular formulas across the library: **Figure 1h** includes only Δm compositions consisting exclusively of C, H, and O, whereas **Figure 1i** includes all remaining formulas that contain additional heteroatoms. All processing and visualizations for Figure 1 were performed in Python (version 3.13.7) using standard scientific libraries, including *pandas*, *matplotlib* and *upsetplot*, and the corresponding scripts are available at https://github.com/helenamrusso/Carnitines_library_manuscript.

### Annotating the molecular formulas of the carnitine modifications

The molecular formulas of the spectra from the carnitine library were predicted through both SIRIUS (version 6.2.2)^50^ and BUDDY (version 1.3).^51^ For BUDDY, the bottom-up MS/MS interrogation was selected, and the experiment-specific global peak annotation was not considered. MS1 and MS/MS tolerance were set to 10 and 20 ppm, respectively. Meta-scores were included for annotation, and adduct forms were defaulted to [M+H]^+^. Chemical element counts were limited as C7-80, H10-150, N1-N6, O3-O20, P0-4, S0-2, F0-4, Cl0-4, Br0-4, I0-4. All the other settings were set as the default.

For SIRIUS, the precursor and fragment ion tolerances were set at 10 and 20 ppm, respectively, specifying Orbitrap profile and considering [M+H]^+^ as ion form. The chemical element counts were defined as: H10-,C7-,N1-,O3-,F0-,P0-,I0-, and the presence of Cl, Br, and S was set to auto detection. The top 10 molecular formulas were calculated for each MS/MS spectrum and were re-ranked using Zodiac^54^ by setting a threshold of 0.95. Molecular fingerprints were generated via CSI:FingerID^55^ by selecting the default databases. The molecular formulas obtained in this final step were considered for determining the molecular formulas of the candidate carnitines.

From the predicted molecular formulas, the formulas of the delta masses were obtained using the *chemparse* (https://pypi.org/project/chemparse/) package in Python. These formulas were added as potential annotations of the modified carnitines in the library.

The resulting resource, GNPS-CANDIDATE-CARNITINES-MASSQL, is now incorporated into the public GNPS propagated spectral library collection (https://gnps.ucsd.edu/ProteoSAFe/gnpslibrary.jsp?library=GNPS-CANDIDATE-CARNITINES-MASSQL).

### Repository-scale searches

The spectra comprising the created carnitines library were searched against public repositories using fast MASST (FASST),^22^ an optimized and accelerated version of the original Mass Spectrometry Search Tool (MASST).^23^ FASST searches are based on cosine similarity between the query spectra and those in the repositories, offering a complementary approach to MassQL-based queries. While MassQL relies on the presence of specific diagnostic ions, instrument-dependent factors, such as limited *m/z* ranges, can result in true carnitine spectra failing to meet strict MassQL criteria. For instance, some datasets do not capture signals at *m/z* 60.0808, a key diagnostic ion used in our queries; however, such spectra may still yield high-confidence matches via cosine similarity.

FASST searches were performed using the RESTful web API (https://zenodo.org/records/7828220) in September 2025, with the *metabolomicspanrepo_index_nightly* database. Search parameters included a cosine similarity threshold of 0.7, a minimum of four matched fragment ions, and precursor and fragment ion tolerances of 0.02 Da. In addition to global spectral matches, domain-specific MASST searches were conducted. plantMASST matches query spectra against 19,075 LC-MS/MS files from taxonomically defined plant extracts.^56^ foodMASST compares results against approximately 3,500 LC-MS/MS files from foods and beverages, organized within a structured food ontology and generated as part of the Global FoodOmics Project.^39,57^ These domain-specific MASSTs enable the mapping of spectral matches to specific files annotated with curated taxonomic or ontological metadata. Collectively, they allow for the identification of the biological or dietary origins of carnitines previously detected in datasets deposited in major metabolomics repositories.

FASST searches are performed on pre-filtered, pre-indexed MS/MS spectra from publicly available datasets, allowing for the searches to be done within seconds. To increase confidence in the spectral matches identified through FASST, a secondary cosine similarity calculation was performed using the original, unfiltered spectra. For this purpose, the MS/MS spectra were retrieved using the Metabolomics Spectrum Resolver API,^58^ and cosine similarity scores were computed using the *matchms* Python package,^59^ and the script for this step is available at https://github.com/helenamrusso/cosine_calculation_FASST_search. These unfiltered cosine scores were merged with the original FASST results, and only matches with cosine similarity scores ≥ 0.7 were retained for downstream analysis.

After applying this filtering step, we obtained matches to 3,872,050 spectra from 220,793 files across 1,977 datasets. To further explore the biological context of these findings, the FASST results were integrated with ReDU (retrieved on September 30, 2025), a controlled vocabulary framework that facilitates systematic comparisons across datasets based on standardized metadata.^24,25^ Datasets associated with humans were identified by the presence of “9606|Homo sapiens” in the NCBITaxonomy column, while rodent datasets were identified using the following taxonomic identifiers: “10088|Mus”, “10090|Mus musculus”, “10105|Mus minutoides”, “10114|Rattus”, and “10116|Rattus norvegicus”. The FASST results were uploaded to https://www.doi.org/10.5281/zenodo.19228538.

### Reanalysis of public datasets from GNPS/MassIVE

To evaluate how the acylcarnitine spectral library can be applied to existing LC-MS/MS data, we selected five publicly available datasets in the GNPS/MassIVE repository that together span different species, matrices, and experimental designs: (1) a mouse study involving antibiotic treatment in colorectal cancer context (MSV000080918); (2) a gnotobiotic mouse experiment comparing germ-free, specific pathogen-free, and monocolonized animals (MSV000088040); (3) a time-resolved mouse feeding experiment with multi-organ and biofluids sampling (MSV000098103); (4) a cohort of mother-infant dyads from Bangladesh (MSV000096943); (5) a large untargeted urine metabolomics dataset comprising about 600 samples from adult women (MSV000096359). For all studies, raw LC-MS/MS files were processed in MZmine 3 or 4^60^, and the parameters used for each study are available in **Supplementary Table S2**, with MZmine batch files available at https://github.com/helenamrusso/Carnitines_library_manuscript. The output files (quantification table and spectral file) were used as input in the Feature-Based Molecular Networking^49^ workflow in the GNPS2 ecosystem^19^, with the acylcarnitine spectral library generated in this study used for spectral matching. Parameters used for the molecular networking and library matching for each use case can be found in **Supplementary Table S3**. The GNPS2 jobs can be accessed through the following links:

- Impact of antibiotic treatment in colorectal cancer dataset (MSV000080918)^61^: https://gnps2.org/status?task=17cb95d221c24806936d1d08fbee6c62
- Monocolonized germ-free mice dataset (MSV000088040)^62,63^: https://gnps2.org/status?task=77658bac1a97469c81c3ffa7f2a74103
- Time-restricted feeding dataset (MSV000098103)^40^: https://gnps2.org/status?task=345d29336f5a4a70b2eeaefae56f8592
- Mother-infant dyads dataset (MSV000096943)^42^: https://gnps2.org/status?task=a6209b0949804988802bfdba3a5f5264
- PlusRise Urobiome dataset (MSV000096359): https://gnps2.org/status?task=8be3e6be2924474db9f24be9252dbb54 (carnitine library) and https://gnps2.org/status?task=b9dfd7e7c57744f8be10955ac6111a83 (all other public libraries)

All the matches obtained with the acylcarnitine library were further filtered by using the PostMN MassQL web application^41^ considering the query below, in which the ion at *m/z* 85.0284 intensity was set to 5% to account for variations among different acquisition methods. Only spectra with library matches that also fulfilled the following MassQL criteria were considered for further analyses.

*QUERY scaninfo(MS2DATA) WHERE MS2PROD=60.0808:TOLERANCEPPM=20:INTENSITYPERCENT=1 AND MS2PROD=85.0284:TOLERANCEPPM=20:INTENSITYPERCENT=5 AND MS2PROD=144.1019:TOLERANCEPPM=20:INTENSITYPERCENT=1*

In addition, the carnitine analogs were assigned structure attributes according to their putative annotation, including acyl chain length (classified as short [C2-C6], medium [C7-C12], long [C13-C21], or very long [C22-C30]), degree of unsaturation, hydroxylation status, and even/odd carbon number.

#### Impact of antibiotic treatment in colorectal cancer dataset

The exported peak areas of the features annotated as carnitines and that also fulfilled MassQL query were used to plot boxplots using the ″seaborn.boxplot” package (version 0.12.2) in Python (version 3.7.6). Significance between the group treated with antibiotic *versus* control was tested using the non-parametric Mann-Whitney U test for antibiotics exposure followed by Benjamini-Hochberg correction (**Supplementary Figure S3a**). The statistical tests were done with the “scipy.stats” package (version 1.7.3), and the p-values were corrected with the “statsmodels.stats.multitest” (version 0.11.1) in Python (version 3.7.6).

#### Monocolonized germ-free mice dataset

For the investigation of this dataset, a heatmap was generated to visualize the log2 fold change of acylcarnitines levels in colonized and monocolonized mice relative to the germ-free group (**Supplementary Figure S3b**). For each feature annotated as an acylcarnitine, the median peak area was calculated per group, and log2(FC) values were computed relative to the germ-free control. Small intestine and colon samples were analyzed separately. The heatmap was constructed using the “seaborn.clustermap” function (seaborn version 0.12.2) in Python (version 3.13.7). Microbial strains were arranged in taxonomic order according to NCBI Taxonomy identifiers, with their corresponding taxonomic classes annotated as a color bar. Acylcarnitines were organized by ascending precursor ion mass, and features with putative CHO-only modifications were annotated using LIPID MAPS^32^ standard nomenclature for carnitines.

#### Time-restricted feeding dataset

The peak areas relative to the carnitine analogs annotated through our library and that fulfilled MassQL query were used for the analyses. Peak area medians were calculated for each feature in each tissue-time point combination. Relative abundances were then calculated by normalizing each metabolite’s abundance to its maximum observed value within each tissue. Stacked bar plots were created with summed abundances grouped by tissue-time point combination using pandas and Matplotlib. Temporal abundance trends across circadian time points were visualized for each tissue using Gaussian-smoothed tendency lines generated by cubic spline interpolation (scipy.interpolate.interp1d) with 1D Gaussian filtering (scipy.ndimage.gaussian_filter1d), with 95% confidence intervals derived from standard error of the aggregated values (**Figure 3a**, **Supplementary Figure 4**).

To identify acyl carnitines with synchronized circadian abundance profiles, pairwise Pearson correlations were computed between all metabolites detected within each tissue or biofluid using relative abundance values across the different timepoints. Only metabolite pairs with non-missing values at a minimum of three time points and a Pearson correlation coefficient ≥ 0.95 were retained. The resulting metabolite-tissue group co-occurrence was visualized as a binary clustermap using seaborn.clustermap function, where rows represent individual carnitine species and columns represent tissue-specific synchronized groups. Filled cells indicate the presence of a given carnitine within that group. Rows and columns were hierarchically clustered using Seaborn’s “sns.clustermap” function with default linkage parameters (average linkage and Euclidean distance). Row color annotations indicate acyl chain length, degree of unsaturation, hydroxylation status, and carbon chain parity for each metabolite (**Figure 3b**). For specific synchronized groups identified through the correlation analysis, individual carnitine relative abundance profiles were visualized as time-course line plots across the time points. For each metabolite, mean relative abundance was calculated across biological replicates for each time point, and curves were smoothed using B-spline interpolation (scipy.interpolate.make_interp_spline) when at least four time points with valid values were available. Plots were generated using Matplotlib and Seaborn, with each metabolite representing a different color (**Figure 3c**).

#### Mother-infant dyads dataset

A public metabolomics dataset from the SEPSiS (Synbiotics for the Early Prevention of Severe Infections in Infants) Observational Cohort Study (MSV000096943) was reanalyzed. A detailed description of the study cohort, sample collection, untargeted metabolomics sample preparation and data acquisition has been reported previously^42^. The feature quantification table and annotation table were imported in R v 4.4.2 (R Foundation for Statistical Computing, Vienna, Austria) for downstream analysis. Blank subtraction was performed by removing features detected in Blank samples when peak areas were not at least 5 times that observed in the samples. Features with RT < 0.2 or > 8 min were excluded, in addition to 3 fecal samples with missing metadata, clustering with plasma samples, and high intensity in the total ion chromatogram. Polymers detected in the LC-MS/MS run were removed using the package ‘homologueDiscoverer v 0.0.0.9000’^64^. Features with near zero variance were removed using the package ‘caret v 6.0’. The ‘lmerTest v 3.1’ package was used to generate the linear mixed effect models. Subject ID was used as random effect and infant age as fixed effect (X ∼ infant_age_days + (1|part_id)). Upset plots were generated using the package ‘UpSetR v 1.4’^65^. For visualization of carnitine trajectories over time, zero values were excluded. Smoothed trend lines were generated using LOESS (non-parametric local regression smoothing method) and added for visualization only.

#### PLUS Consortium RISE for Health dataset

The methods of the IRB-approved, observational RISE study (UMN IRB #00012315) and its biospecimen substudy have been previously described. Metabolomic analyses of these biospecimens were approved by the PLUS Consortium and this secondary analysis of deidentified biospecimens received a not human subjects research designation by the University of California, San Diego Institutional Review Board (IRB #809432). Urine samples were prepared as previously described^66^. The molecular networking obtained for this study was imported to Cytoscape (version 3.10.4) for visualization. Matches to the acyl carnitine library were mapped into the network, in addition to information of scan numbers retrieved in MassQL. Peak areas of the features annotated as carnitine-dihydroferulic acid and carnitine-dihydroferulic acid-sulfate were used to create boxplots using the ″seaborn.boxplot” package (version 0.12.2) in Python (version 3.7.6). Significance between the Body Mass Index (BMI) groups was tested using the non-parametric Mann-Whitney U test followed by Benjamini-Hochberg correction (**Figure 5d**). The statistical tests were done with the “scipy.stats” package (version 1.7.3), and the p-values were corrected with the “statsmodels.stats.multitest” (version 0.11.1) in Python (version 3.7.6). A heatmap showing the log peak areas of carnitine, ferulic acid, and carnitine-dihydroferulic acid and carnitine-dihydroferulic acid-sulfate among all the participants in the study was created using Seaborn (sns.clustermap), with samples (columns) hierarchically clustered using Bray-Curtis dissimilarity (**Figure 5h**).

### Reverse cosine searches

Reverse cosine analysis was applied to obtain putative annotations for unknown carnitine conjugates. Specifically, all candidate carnitine MS/MS spectra, compiled in MGF format, were searched against the GNPS spectral library (ALL_GNPS_NO_PROPOGATED, https://external.gnps2.org/gnpslibrary) using reverse cosine scoring. Matches were retained using the following thresholds: minimum reverse cosine score of 0.7 and at least 3 matched fragment peaks. Computations were performed on a Linux (Ubuntu 20.04) workstation using Python 3.10. The implementation of the reverse cosine algorithm is available at https://github.com/Philipbear/conjugated_metabolome.

### Acyl carnitine synthesis

#### Dihydroferulic acid 4-O-sulfate and 4-O-sulfodihydroferuloylcarnitine

The synthesis performed in this work followed methodologies previously described for multiplexed synthesis^11,13^. Dihydroferulic acid (1 eq.) was dissolved in pyridine (2 mL) in a 20 mL scintillation vial equipped with a magnetic stir bar. Sulfur trioxide–pyridine complex (1.5 eq.) was added portion wise, and the reaction mixture was stirred at 23 °C overnight. The mixture was concentrated in vacuo and used directly in the subsequent step.

Dihydroferulic acid 4-*O*-sulfate (1 eq.) was dissolved in DMF (2 mL) in a 20 mL scintillation vial equipped with a magnetic stir bar. Solid EDC·HCl (1.2 eq.) and DMAP (0.1 eq.) were added sequentially, and the mixture was stirred at 23 °C. After 15 min, carnitine (1.2 eq.) was added, and the reaction was stirred for 14 h. The mixture was concentrated under reduced pressure, and the residue was dissolved in an appropriate solvent and filtered prior to LC–MS/MS analysis.

#### Dihydroferuloylcarnitine purification

Dihydroferulic acid (1 eq.) was dissolved in DMF (2 mL) in a 20 mL scintillation vial equipped with a magnetic stir bar. EEDQ (1.2 eq.) and carnitine (5 eq.) were added sequentially, and the reaction was stirred at room temperature for 14 h. The mixture was concentrated under reduced pressure, and the residue was purified using a CombiFlash NextGen 300+ with reversed phase column C18 100 g Gold at a flow rate 10 mL per min with H_2_O (Solvent A) and ACN (solvent B) with the following gradient: 0-5 min, 0% B; 5-14 min, 1% B; 14-20 min 40% B; 20-25 min, 80% B. Dihydroferuloylcarnitine eluted at 10 min, 1% B. ^1^H-NMR (D_2_O) δ 6.81 (s, 1H), 6.72 (d, 1H), 6.62 (d, 1H), 5.43 (m, 1H), 3.73 (s, 3H), 3.61-3.47 (dd, 2H), 2.86 (s, 9H), 2.76 (t, 2H), 2.67(m, 2H), 2.58 (t, 2H). 13C NMR (151 MHz, Deuterium Oxide) δ 174.2, 172.7, 147.3, 143.3, 132.9, 121.2, 115.4, 112.7, 67.4, 65.1, 55.8, 53.5, 36.6, 35.5, 29.6. The spectra are available at https://doi.org/10.5281/zenodo.19115214.

NMR spectra were collected at 298 K on a 500 MHz Bruker Avance III spectrometer fitted with a 1.7 mm triple resonance cryoprobe with z-axis gradients. ^1^H NMR: D_2_O (4.7), CDCl_3_ (7.26) at 500 MHz. Dihydroferuloylcarnitine spectra was taken in D_2_O with shifts reported in parts per million (ppm) referenced to the proton of the solvent (4.7). Data for ^1^H-NMR are reported as follows: chemical shift (ppm, reference to protium; s = single, d = doublet, t = triplet, q = quartet, dd = doublet of doublets, m = multiplet, coupling constant (Hz), and integration).

### Retention time and drift time analysis

Both the biological sample (P4_A2_364302537) and the reaction mixtures were injected (3 µL) into a Vanquish UHPLC system coupled to a Q Exactive Orbitrap mass spectrometer (Thermo Fisher Scientific). Initially, the chromatographic separation was achieved by reverse-phase polar C18 (Kinetex Polar C18, 100 × 2.1 mm, 2.6 μm particle size, 100 A pore size; Phenomenex, Torrance) at 40 °C column temperature. The mobile phase consisted of solvents A (water) and B (ACN), both containing 0.1% formic acid, and the flow rate was set at 0.5 mL/min. The chromatographic gradient employed was the following: 0-0.5 min 5% B, 0.5-8.0 min 5-99% B, 8.0- 10.0 min 99% B, 10.0-10.2 min 99-5% B, 10.2-12.2 min 5% B. Another chromatographic condition was used employing a HILIC ACQUITY^TM^ Premier column (Glycan BEH Amide, 100 × 2.1 mm, 1.7 µm particle size, 130 A pore size). at 40 °C column temperature. The mobile phase consisted of solvents A (50 mM ammonium formate, pH 4.4) and B (ACN), and the flow rate was set at 0.5 mL/min. The chromatographic gradient employed was the following: 0-0.5 min 95% B, 0.5-8.0 min 95-5% B, 8.0-10.0 min 5% B, 10.0-10.2 min 5-95% B, 10.2-12.2 min 95% B. Mass spectrometry (MS) analysis was performed using electrospray ionization (ESI) in positive ionization mode, and the parameters were set as: sheath gas flow 53 L/min, auxiliary gas flow rate 14 L/min, sweep gas flow 3 L/min, spray voltage 3.5 kV, inlet capillary to 269°C, and auxiliary gas heater 430 °C. MS1 scan range was set to *m/z* 100-1,000 with a resolution (R*_m/z_* _200_) of 35,000, automatic gain control (AGC) target as 5.0E4, and maximum injection time of 100 ms. Up to 5 MS/MS spectra per MS1 were collected with a resolution (R*_m/z_* _200_) set to 17,500, AGC target as 1.0E5, and maximum injection time of 100 ms. The isolation window was set to 1.0 *m/z* and the isolation offset was set to 0 m/z. The normalized collision energy was acquired with an increased stepwise from 25 to 40 to 60%. The apex trigger was set to 1 to 5 s, the minimum AGC target for the MS/MS spectrum was 8.0E3, and a dynamic precursor exclusion of 10 s was selected.

For drift time analysis, a timsMetabo mass spectrometer (Bruker Daltonics) with the VIP-HESI source coupled to an Agilent 1260 Infinity 3 UHPLC was employed. Both the reaction mixtures and biological sample were injected (3 µL) to the liquid chromatography system equipped with a reverse-phase polar C18 (Kinetex Polar C18, 100 × 2.1 mm, 2.6 μm particle size, 100 A pore size; Phenomenex, Torrance) at 40 °C column temperature. The mobile phase consisted of solvents A (water) and B (ACN), both containing 0.1% formic acid, and the flow rate was set at 0.5 mL/min. The chromatographic gradient employed was the following: 0-0.5 min 5% B, 0.5-5.0 min 5-15% B, 5.0-5.1 min 99% B, 5.1-8.1 min 99% B, 8.1-8.2 min 99-5% B, 8.2-10.2 min 5% B. The instrument was operated in positive ionization mode with capillary voltage 3500 V, end-plate offset 500 V, nebulizer pressure 0.2 bar, dry gas 8.0 L/min, and dry gas temperature 200 °C. MS data were acquired from m/z 20–1300 in PASEF mode. TIMS mobility was collected with a 1/K0 range of 0.30–1.50 V·s/cm2, ramp time of 100 ms. The Athena Ion Processor (AIP) was turned on to 600-0Vpp_slope-20Vpp/us for MS and MS/MS. External mass and mobility calibration were performed using the Bruker ESI Tuning Mix (ESI-TOF CCS compendium). Extracted ion mobilograms and chromatograms of the [M+H]+ of each targeted ion were exported through the Bruker DataAnalysis 6.1 software. Raw data was converted to open-source “.mzML” format in TIMSCONVERT^67^ through centroiding, zlib compression, 64-bit encoding, and combining all spectra into a single file.

## Code Availability

All the scripts used to perform the data analyses and generate the figures are available at https://github.com/helenamrusso/Carnitines_library_manuscript and https://github.com/kinekvitne/manuscript_carnitines_library. Scripts used for the library generation from MassQL results are available at https://github.com/helenamrusso/library_generation_from_MassQL.

## Data Availability

All the untargeted metabolomics LC-MS/MS data used or generated in this study are deposited on GNPS/MassIVE and are publicly available under the following accession numbers: MSV000080918 (antibiotics treatment), MSV000088040 (monocolonized germ-free mice), MSV000098103 (time-restricted feeding dataset), MSV000096943 (mother-infant dyads dataset), MSV000098103 (PlusRise Urobiome), MSV000101216 (retention time and drift time analysis). NMR analysis of carnitine-dihydroferulic acid is archived at Zenodo (https://doi.org/10.5281/zenodo.19115214). The acyl carnitine library is available as part of the GNPS public spectral libraries (https://gnps.ucsd.edu/ProteoSAFe/gnpslibrary.jsp?library=GNPS-CANDIDATE-CARNITINES-MASSQL). The FASST results can be found at https://www.doi.org/10.5281/zenodo.19228538.

## References

1. Dambrova, M. et al. Acylcarnitines: Nomenclature, biomarkers, therapeutic potential, drug targets, and clinical trials. Pharmacol. Rev. 74, 506–551 (2022).

2. Miller, M. J., Cusmano-Ozog, K., Oglesbee, D., Young, S. & ACMG Laboratory Quality Assurance Committee. Laboratory analysis of acylcarnitines, 2020 update: a technical standard of the American College of Medical Genetics and Genomics (ACMG). Genet. Med. 23, 249–258 (2021).

3. Violante, S. et al. Substrate specificity of human carnitine acetyltransferase: Implications for fatty acid and branched-chain amino acid metabolism. Biochim. Biophys. Acta 1832, 773–779 (2013).

4. Gotvaldová, K. et al. BCAA metabolism in pancreatic cancer affects lipid balance by regulating fatty acid import into mitochondria. Cancer Metab. 12, 10 (2024).

5. Benvenga, S., Lakshmanan, M. & Trimarchi, F. Carnitine is a naturally occurring inhibitor of thyroid hormone nuclear uptake. Thyroid 10, 1043–1050 (2000).

6. Sinclair, C., Gilchrist, J. M., Hennessey, J. V. & Kandula, M. Muscle carnitine in hypo- and hyperthyroidism. Muscle Nerve 32, 357–359 (2005).

7. Tortorelli, S. et al. The urinary excretion of glutarylcarnitine is an informative tool in the biochemical diagnosis of glutaric acidemia type I. Mol. Genet. Metab. 84, 137–143 (2005).

8. Arjmand, B. et al. Association of plasma acylcarnitines and amino acids with hypertension: A nationwide metabolomics study. PLoS One 18, e0279835 (2023).

9. Vianey-Saban, C., Fouilhoux, A., Vockley, J., Acquaviva-Bourdain, C. & Guffon, N. Improving diagnosis of mitochondrial fatty-acid oxidation disorders. Eur. J. Hum. Genet. 31, 265–272 (2023).

10. Chatterjee, P. et al. Plasma metabolites associated with biomarker evidence of neurodegeneration in cognitively normal older adults. J. Neurochem. 159, 389–402 (2021).

11. Patan, A. et al. Charting the undiscovered metabolome with synthetic multiplexing. bioRxiv (2025) doi:10.1101/2025.11.18.689170.

12. Charron-Lamoureux, V. et al. A guide to reverse metabolomics-a framework for big data discovery strategy. Nat. Protoc. (2025) doi:10.1038/s41596-024-01136-2.

13. Gentry, E. C. et al. Reverse metabolomics for the discovery of chemical structures from humans. Nature (2023) doi:10.1038/s41586-023-06906-8.

14. Yan, X. et al. Mass spectral library of acylcarnitines derived from human urine. Anal. Chem. 92, 6521–6528 (2020).

15. Damiani, T. et al. A universal language for finding mass spectrometry data patterns. Nat. Methods 22, 1247–1254 (2025).

16. Mannochio-Russo, H. et al. The microbiome diversifies long- to short-chain fatty acid-derived N-acyl lipids. Cell (2025) doi:10.1016/j.cell.2025.05.015.

17. Mohanty, I. et al. The Underappreciated Diversity of Bile Acid Modifications. (2023) doi:10.2139/ssrn.4436846.

18. Patterson, A. et al. Glucuronidation metabolomic fingerprinting to map host-microbe metabolism. Res. Sq. (2025) doi:10.21203/rs.3.rs-6321321/v1.

19. Wang, M. et al. Sharing and community curation of mass spectrometry data with Global Natural Products Social Molecular Networking. Nat. Biotechnol. 34, 828–837 (2016).

20. Yurekten, O. et al. MetaboLights: open data repository for metabolomics. Nucleic Acids Res. 52, D640–D646 (2024).

21. Sud, M. et al. Metabolomics Workbench: An international repository for metabolomics data and metadata, metabolite standards, protocols, tutorials and training, and analysis tools. Nucleic Acids Res. 44, D463–70 (2016).

22. Batsoyol, N., Pullman, B., Wang, M., Bandeira, N. & Swanson, S. P-Massive: A Real-Time Search Engine for a Multi-Terabyte Mass Spectrometry Database. in SC22: International Conference for High Performance Computing, Networking, Storage and Analysis 1–15 (2022).

23. Wang, M. et al. Mass spectrometry searches using MASST. Nat. Biotechnol. 38, 23–26 (2020).

24. El Abiead, Y., et al. Enabling pan-repository reanalysis for big data science of public metabolomics data. ChemRxiv (2024) doi:10.26434/chemrxiv-2024-jt46s.

25. Jarmusch, A. K. et al. ReDU: a framework to find and reanalyze public mass spectrometry data. Nat. Methods 17, 901–904 (2020).

26. Wishart, D. S. et al. HMDB 5.0: the Human Metabolome Database for 2022. Nucleic Acids Res. 50, D622–D631 (2022).

27. Kim, S. et al. PubChem 2023 update. Nucleic Acids Res. 51, D1373–D1380 (2023).

28. Zhang, J. et al. Simultaneously quantifying hundreds of acylcarnitines in multiple biological matrices within ten minutes using ultrahigh-performance liquid-chromatography and tandem mass spectrometry. J. Pharm. Anal. 14, 140–148 (2024).

29. Keys, A. J. et al. Exploring acylcarnitine metabolism using reverse metabolomics. ChemRxiv (2026) doi:10.26434/chemrxiv.15000426/v1.

30. Bittremieux, W., Laukens, K., Noble, W. S. & Dorrestein, P. C. Large-scale tandem mass spectrum clustering using fast nearest neighbor searching. Rapid Commun. Mass Spectrom. 39 Suppl 1, e9153 (2025).

31. Watrous, J. et al. Mass spectral molecular networking of living microbial colonies. Proc. Natl. Acad. Sci. U. S. A. 109, E1743–52 (2012).

32. Conroy, M. J. et al. LIPID MAPS: update to databases and tools for the lipidomics community. Nucleic Acids Res. 52, D1677–D1682 (2024).

33. Song, X. et al. Dark reactions in microdroplets explain widespread artifacts in metabolomic profiling. ACS Meas. Sci. Au (2026) doi:10.1021/acsmeasuresciau.5c00146.

34. Schönfeld, P. & Wojtczak, L. Short- and medium-chain fatty acids in energy metabolism: the cellular perspective. J. Lipid Res. 57, 943–954 (2016).

35. Hastings, J. et al. ChEBI in 2016: Improved services and an expanding collection of metabolites. Nucleic Acids Res. 44, D1214–9 (2016).

36. Jacques, F., Rippa, S. & Perrin, Y. Physiology of L-carnitine in plants in light of the knowledge in animals and microorganisms. Plant Sci. 274, 432–440 (2018).

37. Zuffa, S., et al. Influence of perinatal ampicillin exposure on maternal fecal microbial and metabolic profiles. bioRxivorg (2025) doi:10.1101/2025.06.30.662372.

38. Zuffa, S., et al. A multi-organ Murine metabolomics atlas reveals molecular dysregulations in Alzheimer’s Disease. bioRxiv (2025) doi:10.1101/2025.04.28.651123.

39. West, K. A., Schmid, R., Gauglitz, J. M., Wang, M. & Dorrestein, P. C. foodMASST a mass spectrometry search tool for foods and beverages. NPJ Sci Food 6, 22 (2022).

40. Reilly, E. R. et al. Systemic rhythmicity of host and bacterial bile acid amidates in the mouse. Cell Syst. 101541 (2026).

41. Mannochio-Russo, H. et al. Bridging complexity and accessibility in metabolomics with MetaboApps. ChemRxiv (2025) doi:10.26434/chemrxiv-2025-3nq29.

42. Kvitne, K. E. et al. Environmental and maternal imprints on infant gut metabolic development. Cell Host Microbe 33, 2130–2147.e7 (2025).

43. Ouyang, R. et al. Maturation of the gut metabolome during the first year of life in humans. Gut Microbes 15, 2231596 (2023).

44. Smith, A. L. et al. RISE FOR HEALTH: Rationale and protocol for a prospective cohort study of bladder health in women. Neurourol. Urodyn. 42, 998–1010 (2023).

45. Xing, S., et al. Navigating the conjugated metabolome. bioRxivorg (2026) doi:10.64898/2026.02.06.704496.

46. Xing, S. et al. Reverse spectral search reimagined: A simple but overlooked solution for chimeric spectral annotation. Anal. Chem. 97, 17926–17930 (2025).

47. Sumner, L. W. et al. Proposed minimum reporting standards for chemical analysis Chemical Analysis Working Group (CAWG) Metabolomics Standards Initiative (MSI). Metabolomics 3, 211–221 (2007).

48. El Abiead, Y., et al. Structure-centric searching enables global mapping of the public metabolome. Nat. Biotechnol. In press.

49. Nothias, L.-F. et al. Feature-based molecular networking in the GNPS analysis environment. Nat. Methods 17, 905–908 (2020).

50. Dührkop, K. et al. SIRIUS 4: a rapid tool for turning tandem mass spectra into metabolite structure information. Nat. Methods 16, 299–302 (2019).

51. Xing, S., Shen, S., Xu, B., Li, X. & Huan, T. BUDDY: molecular formula discovery via bottom-up MS/MS interrogation. Nat. Methods 20, 881–890 (2023).

52. da Silva, R. R., Dorrestein, P. C. & Quinn, R. A. Illuminating the dark matter in metabolomics. Proceedings of the National Academy of Sciences of the United States of America vol. 112 12549–12550 (2015).

53. El Abiead, Y., et al. A perspective on unintentional fragments and their impact on the dark metabolome, untargeted profiling, molecular networking, public data, and repository scale analysis. JACS Au 5, 5828–5850 (2025).

54. Ludwig, M. et al. Database-independent molecular formula annotation using Gibbs sampling through ZODIAC. Nature Machine Intelligence 2, 629–641 (2020).

55. Dührkop, K., Shen, H., Meusel, M., Rousu, J. & Böcker, S. Searching molecular structure databases with tandem mass spectra using CSI:FingerID. Proc. Natl. Acad. Sci. U. S. A. 112, 12580–12585 (2015).

56. Gomes, P. W. P., et al. plantMASST - Community-driven chemotaxonomic digitization of plants. bioRxiv (2024) doi:10.1101/2024.05.13.593988.

57. Gauglitz, J. M. et al. Enhancing untargeted metabolomics using metadata-based source annotation. Nat. Biotechnol. 40, 1774–1779 (2022).

58. Bittremieux, W. et al. Universal MS/MS Visualization and Retrieval with the Metabolomics Spectrum Resolver Web Service. bioRxiv 2020.05.09.086066 (2020) doi:10.1101/2020.05.09.086066.

59. Huber, F. et al. Matchms - processing and similarity evaluation of mass spectrometry data. J. Open Source Softw. 5, 2411 (2020).

60. Schmid, R. et al. Integrative analysis of multimodal mass spectrometry data in MZmine 3. Nat. Biotechnol. (2023) doi:10.1038/s41587-023-01690-2.

61. Shalapour, S. et al. Inflammation-induced IgA+ cells dismantle anti-liver cancer immunity. Nature 551, 340–345 (2017).

62. Song, X. et al. Gut microbial fatty acid isomerization modulates intraepithelial T cells. Nature 619, 837–843 (2023).

63. Wu, M. et al. Gut complement induced by the microbiota combats pathogens and spares commensals. Cell 187, 897–913.e18 (2024).

64. Mildau, K. et al. Homologue series detection and management in LC-MS data with homologueDiscoverer. Bioinformatics 38, 5139–5140 (2022).

65. Conway, J. R., Lex, A. & Gehlenborg, N. UpSetR: an R package for the visualization of intersecting sets and their properties. Bioinformatics 33, 2938–2940 (2017).

66. Weldon, K. C. et al. Urinary metabolomic profile is minimally impacted by common storage conditions and additives. Int. Urogynecol. J. 36, 839–847 (2025).

67. Luu, G. T. et al. TIMSCONVERT: a workflow to convert trapped ion mobility data to open data formats. Bioinformatics 38, 4046–4047 (2022).

