## Supplementary Figures for "Pan-Metabolomics Repository Mapping of the Carnitine Landscape"

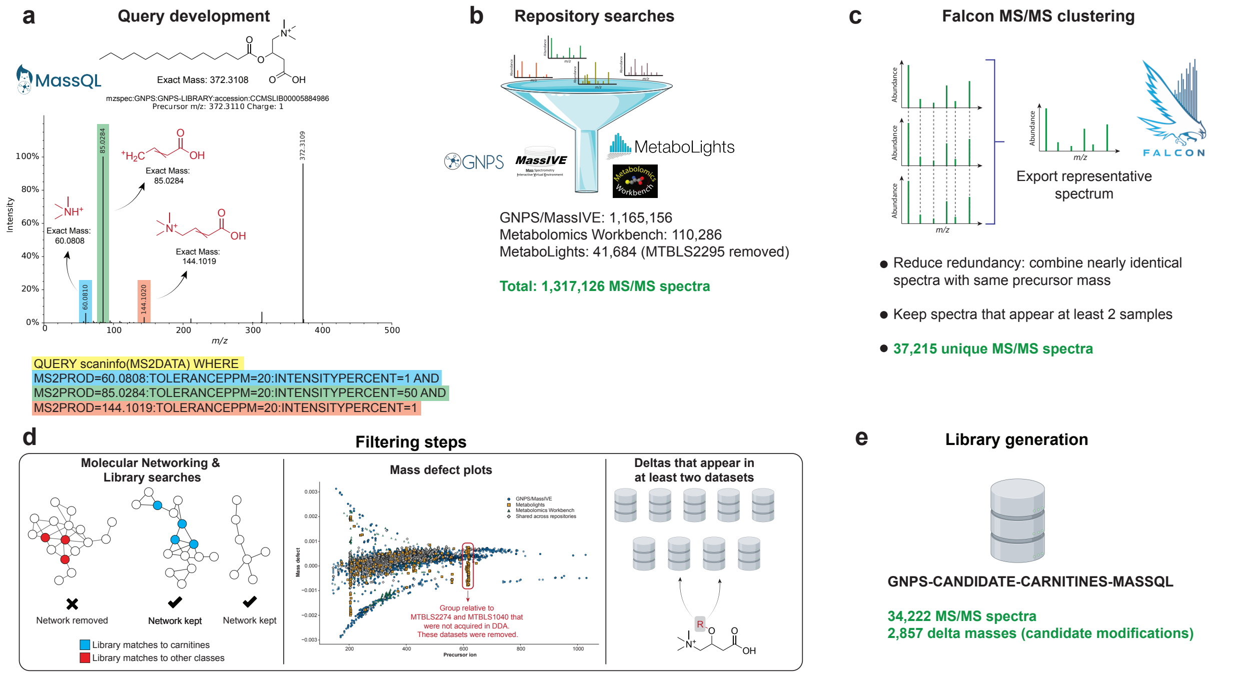


**Supplementary Figure S1. Workflow for MassQL-based retrieval and curation of candidate acylcarnitine MS/MS spectra from public repositories. (a)** Development of a MassQL query in positive ionization mode using three diagnostic fragment ions of acylcarnitines. **(b)** Application of this query to GNPS/MassIVE, MetaboLights and Metabolomics Workbench. **(c)** Falcon MS/MS clustering to reduce redundancy and keep spectra observed in at least two samples. **(d)** Additional curation using molecular networking, mass-defect plots and dataset-level filters to discard non-carnitine networks and aberrant precursor regions, and to retain only delta masses seen in ≥ 2 independent datasets. **(e)** Final GNPS-CANDIDATE-CARNITINES-MASSQL library comprising 34,222 MS/MS spectra and 2,587 unique delta masses (putative acylcarnitine modifications).

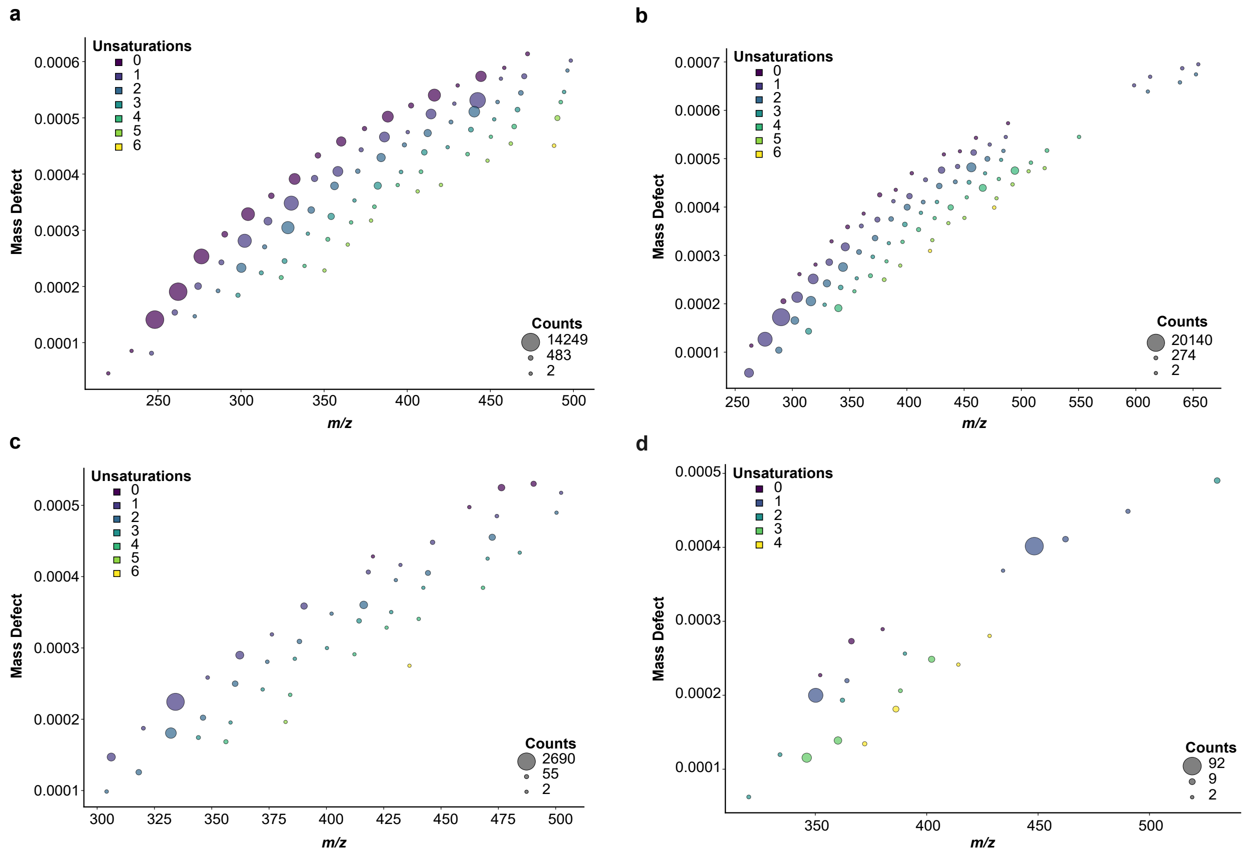


**Supplementary Figure S2. Mass defect plots of CHO candidate acylcarnitines grouped by oxygen content**. **(a-d)** Putative CHO-only acylcarnitine annotations are plotted as precursor *m/z* versus mass defect, with panels corresponding to elemental compositions containing one **(a)**, two **(b)**, three **(c)**, or four **(d)** oxygen atoms. Point size is proportional to the number of detections across all repositories, and point color encodes the number of unsaturations.

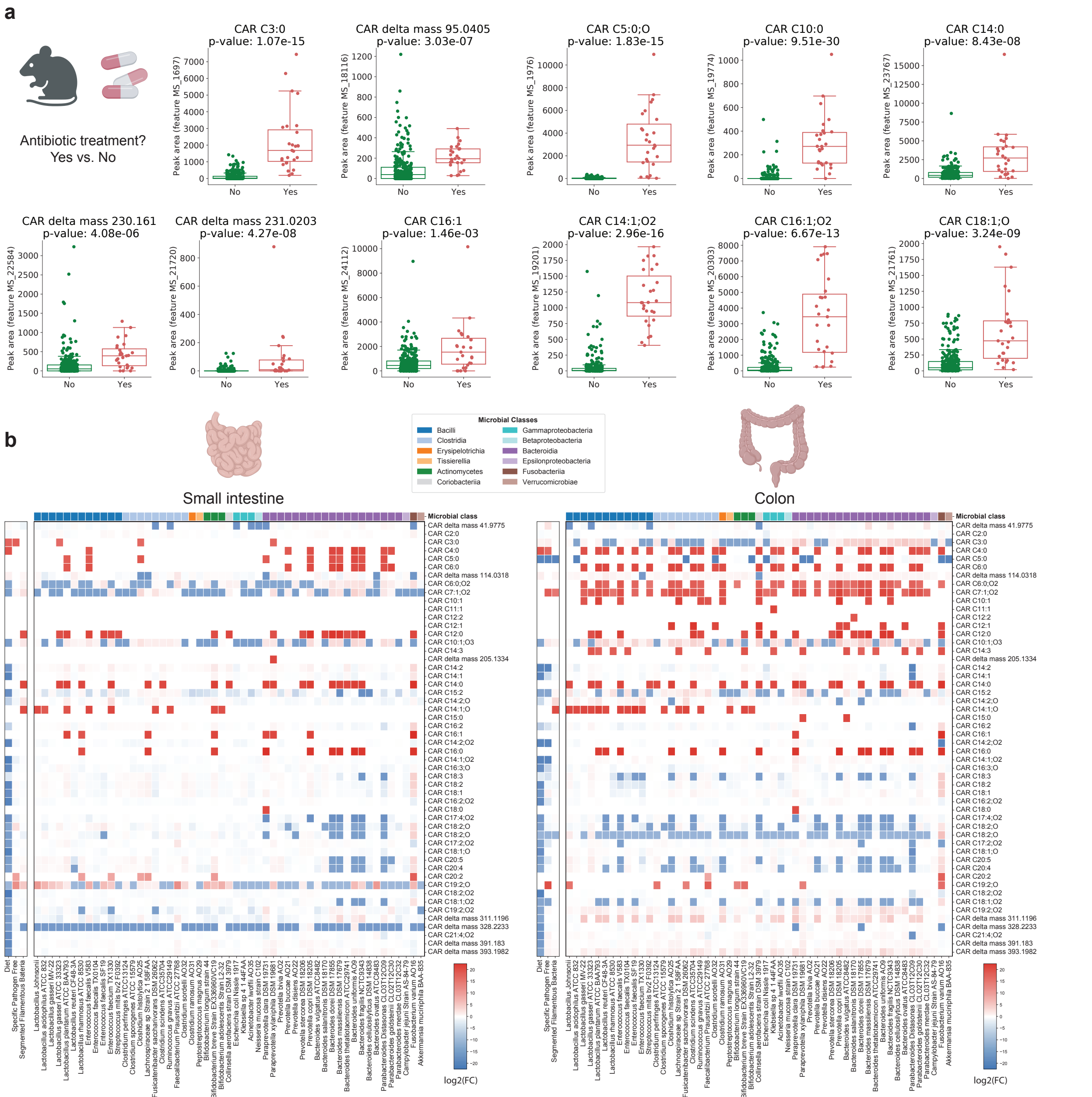


**Supplementary Figure S3. Application of the carnitine library to a monocolonized germ-free mice dataset. (a)** Selected carnitine-related features associated with antibiotic treatment in monocolonized germ-free mice (antibiotics, n = 310; no antibiotics, n = 27; obese, n = 136). Significance was tested using the non-parametric Mann-Whitney U test and p values were corrected for multiple comparisons using the Benjamini-Hochberg correction. Boxplots indicate the first (lower), median, and third (upper) quartiles, while whiskers are 1.5 times the interquartile range. **(b)** Heatmap log2 fold changes (FCs) of the acylcarnitines matches in colon and small intestine samples of monocolonized mice relative to GF mice. Red cells indicate compounds that are increasing relative to GF, while blue cells indicate compounds that are decreasing relative to GF mice. The x axis is taxonomically ordered according to the NCBI Taxonomy ID. Carnitine derivatives showed microbiome-dependent remodeling across monocolonized gut microbes, with taxon-specific changes relative to germ-free controls in the small intestine and colon, with a broader increase in C2-C16 species that was more pronounced in the colon. Icons were obtained from BioRender.

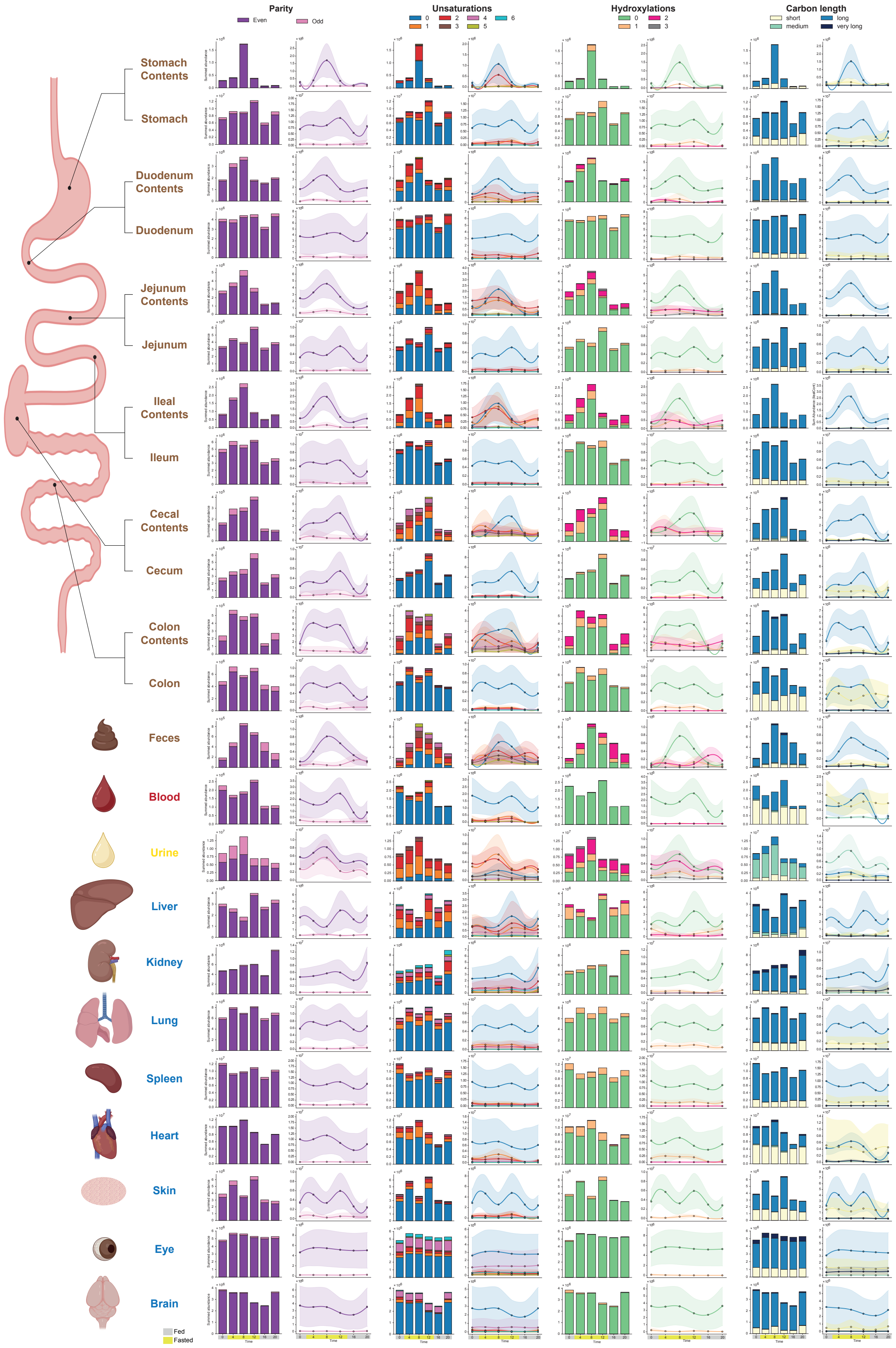


**Supplementary Figure S4. Diurnal patterns of acyl carnitine profiles across 23 mouse tissues and biofluids[1] stratified by structural class.** For each tissue or biofluid (rows), summed peak areas per time point are displayed as stacked bar plots and circadian trends are shown as Gaussian-smoothed tendency lines for four structural classifications: carbon chain parity (Even/Odd), degree of unsaturation (0-6), number of hydroxylations (0-3), and acyl chain length (short, C2-C6; medium, C7-C12; long, C13-C21; very long, C22-C30). Shaded bands indicate 95% confidence intervals calculated from the standard error of the sum. Individual points show the sum of all detected carnitines within each chain length category at each time point. Icons were obtained from BioRender.

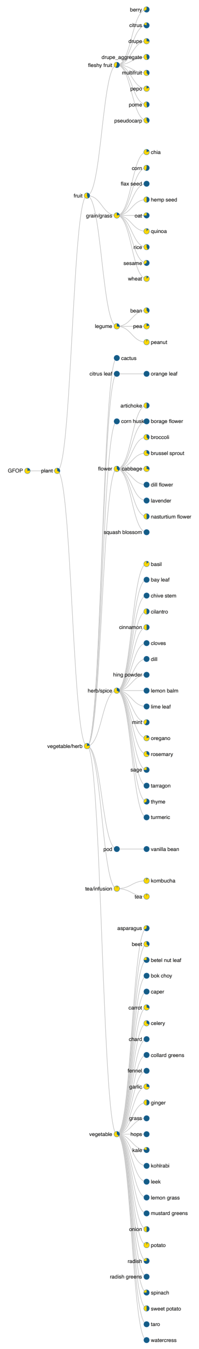


**Supplementary Figure S5. foodMASST results using ferulic acid as input.** The search was done using StructureMASST's FASSTrecords functionality.

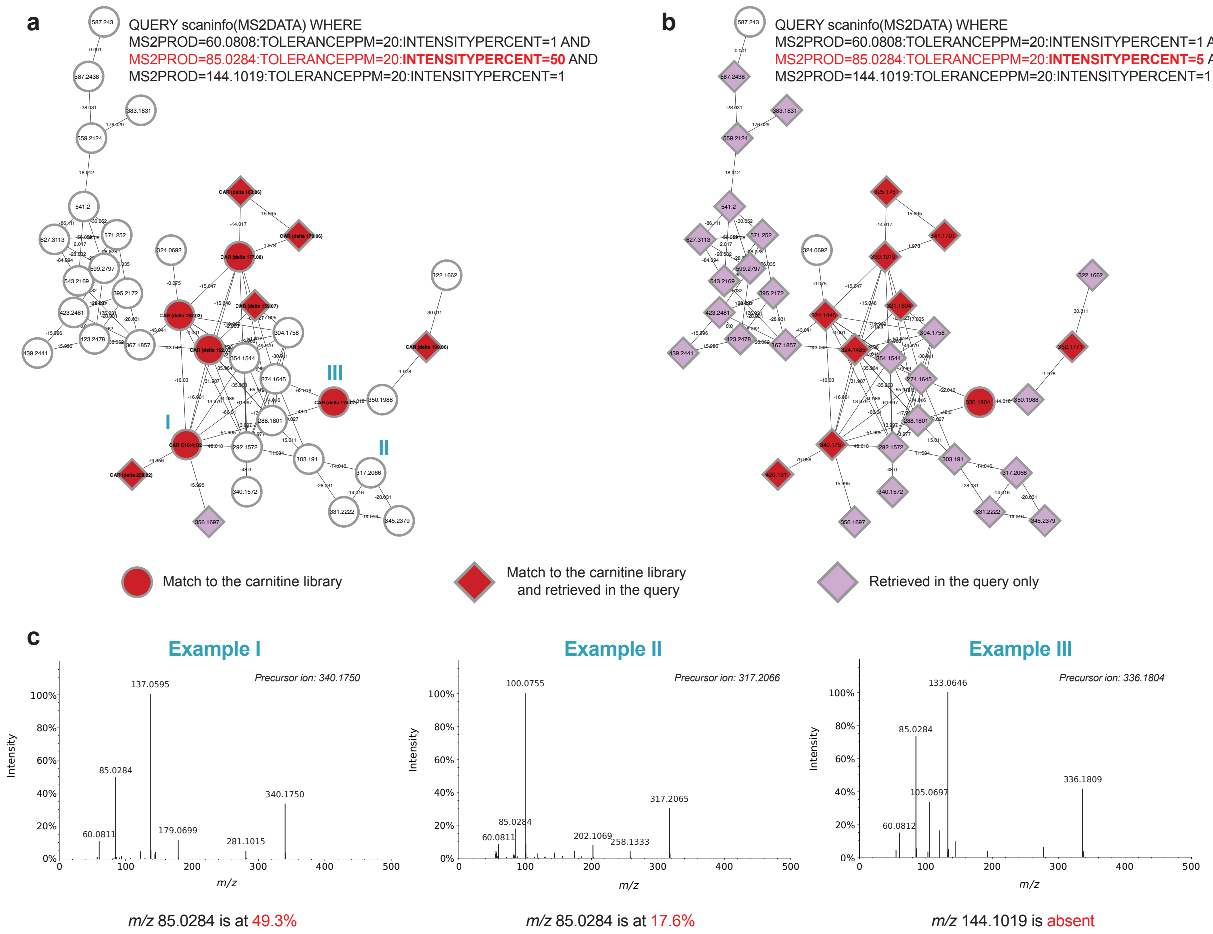


**Supplementary Figure S6. Effect of MassQL intensity threshold on acylcarnitine spectral retrieval.** **(a,b)** Selected subnetwork of the feature-based molecular network[2] of carnitine-related metabolites detected in the PlusRise Urobiome study[3]. Diamond-shaped nodes are relative to spectra retrieved from the queries depicted: **(a)** the original MassQL query with a 50% minimum intensity threshold for the diagnostic ion m/z 85.0284, and **(b)** after lowering the intensity threshold of this ion to 5%, substantially increasing the number of spectra retrieved. **(c)** Representative MS/MS spectra illustrating three scenarios of diagnostic ion behavior across the network. Example **I** shows a spectrum annotated by the carnitine library generated in this work, in which the *m/z* 85.0284 is present at 49.3% relative intensity, falling just below the original 50% MassQL intensity threshold and therefore not retrieved by the original query but recovered upon threshold relaxation to 5%. Example **II** shows a spectrum not annotated by the library in which *m/z* 85.0284 is reduced to 17.6%, retrieved exclusively by the relaxed (5%) MassQL query but not the original (50%) query, illustrating that diagnostic ion intensity progressively decreases for nodes more distant in the network. Example **III** shows a spectrum in which *m/z* 144.1019 is absent, and which is not retrieved by either the original or relaxed MassQL query threshold, neither by the library generated.

**References:**

1. Reilly, E. R. *et al.* Systemic rhythmicity of host and bacterial bile acid amidates in the mouse. *Cell Syst.* 101541 (2026).
2. Nothias, L.-F. *et al.* Feature-based molecular networking in the GNPS analysis environment. *Nat. Methods* **17**, 905–908 (2020).

3. Smith, A. L. *et al.* RISE FOR HEALTH: Rationale and protocol for a prospective cohort study of bladder health in women. *Neurourol. Urodyn.* **42**, 998–1010 (2023).
